# Diminazene resistance in *Trypanosoma congolense* is linked to reduced mitochondrial membrane potential and not to reduced transport capacity

**DOI:** 10.1101/2020.07.28.224543

**Authors:** Lauren V. Carruthers, Jane C. Munday, Godwin U. Ebiloma, Pieter Steketee, Siddharth Jayaraman, Gustavo D. Campagnaro, Marzuq A. Ungogo, Anne-Marie Donachie, Tim G. Rowan, Rose Peter, Liam J. Morrison, Michael P. Barrett, Harry P. De Koning

## Abstract

*Trypanosoma congolense* is one of the principal agents causing livestock trypanosomiasis in sub-Saharan Africa. This wasting disease is costing these developing economies billions of dollars and undermining food security. Only two old drugs, the diamidine diminazene and the phenanthridine isometamidium are regularly used, and resistance is widespread but poorly understood. We induce diminazene resistance in *T. congolense* laboratory strain IL3000 in vitro. Resistance was stable and not deleterious to in vitro growth. There was no cross-resistance with the phenanthridine drugs, with melaminophenyl arsenicals, with two promising new oxaborole trypanocides, nor with other diamidine trypanocides such as pentamidine, except the close structural analogues DB829 and DB75. Fluorescence microscopy showed that accumulation of DB75 was inhibited by folate. Uptake of [^3^H]-diminazene was also partly inhibited by folate, as well as by competing diamidine drugs, albeit at quite high concentrations, and uptake of tritiated diminazene and pentamidine was slow and low affinity. Uptake of [^3^H]-folate was in turn partly inhibited by diminazene, and inhibition of diminazene uptake by folate and pentamidine appeared to be additive, indicating multiple low affinity transport mechanisms for the drug. Expression of the *T. congolense* folate transporters TcoFT1-3 in diminazene-resistant *T. b. brucei* significantly sensitized the cells to diminazene and DB829, but not to oxaborole AN7973. However, [^3^H]-diminazene uptake was not different in *T. congolense* IL3000 and its diminazene resistant clones and RNAseq and whole-genome sequencing of multiple resistant clones did not reveal major changes in folate transporter sequence or expression. Instead, flow cytometry revealed a strong and stable reduction in the mitochondrial membrane potential Ψm in all resistant clones. We conclude that diminazene uptake in *T. congolense* proceed via multiple low affinity mechanisms including folate transporters and that resistance is the result of a reduction in Ψm that limits mitochondrial accumulation of the drug.

## Introduction

Animal African Trypanosomiasis (AAT), also called nagana, is an infection of livestock and wild animal species with African trypanosomes, principally *Trypanosoma congolense*, *T. brucei* sspp, *T. vivax* and *T. suis*, which is spread by the bite of infected tsetse flies. It has a very severe impact on livestock farming in sub-Saharan Africa, costing billions of dollars in lost agricultural production. As such it is has knock-on effects on issues such as food security, diet and rural development, and the livelihood of rural African communities in endemic areas [1]. Although Human African Trypanosomiasis (HAT), or sleeping sickness, is in rapid decline in most of Africa due to active case finding and rapid treatment [2,3], AAT remains a very serious problem throughout the continent. This is due in no small measure because just two old drugs have been the mainstay of AAT treatment for decades: diminazene aceturate (DA) and isometamidium chloride (ISM). Resistance to each has been reported from most countries in which the disease is prevalent [1] and no new drugs have been introduced for over half a century. Yet, while there are many studies documenting treatment failures in the field, relatively little progress has been made in understanding the mechanisms of resistance in the clinically most relevant animal trypanosome species *Trypanosoma congolense* and *T. vivax*.

In contrast, much progress has been made in understanding resistance to the suite of drugs used to treat the various manifestations of HAT, caused by the *T. brucei* subspecies *T. b. gambiense* in West and Central Africa, and *T. b. rhodesiense* in East and Southern Africa. Most of the resistance mechanisms uncovered for HAT drugs involve aspects of drug uptake by the parasite. For instance, pentamidine (used for early stage *gambiense* HAT) and melarsoprol (now mostly used for late stage *rhodesiense* HAT), are internalized by two transport proteins in the trypanosome’s plasma membrane. The first of these is an aminopurine transporter called P2 [4] encoded by the gene *TbAT1* [5], the deletion of which has been shown to cause a loss of sensitivity to these drugs [6]. Subsequently, it was shown a second and more important determinant for pentamidine-melarsoprol cross-resistance is the High Affinity Pentamidine Transporter (HAPT1) [7,8], which was more recently shown to be the aquaglyceroporin TbAQP2 [9,10]. Rearrangements in the TbAQP2 locus were clearly linked to treatment failures, particularly with melarsoprol [11,12]. Similarly, resistance to eflornithine, used against late-stage *gambiense* HAT, is associated with the loss of an amino acid transporter, TbAAT6 [13,14], and resistance to suramin, the first-line drug against early-stage *rhodesiense* HAT, is the result of mutations that interfere with its uptake by receptor-mediated endocytosis [14,15].

In *T. congolense*, resistance to isometamidium has also been attributed to reduced accumulation of the drug [16–19], possibly in part linked to reduced mitochondrial membrane potential [20,21], which diminishes sequestration of cationic drugs into the mitochondrion [22–24] - the likely site of drug action. For DA, it has been shown in the *T. b. brucei* model that it is almost exclusively taken up by the P2/TbAT1 transporter [25] as it is a very poor substrate for the HAPT1/TbAQP2 transporter [26,27]. It was subsequently proposed that the same mechanism would apply in *T. congolense*, with an adenosine transporter designated TcoAT1 mediating DA uptake, and DA resistance being the result of specific single nucleotide polymorphisms (SNPs) that could be used to screen for resistance [28–30]. However, expression of the putative TcoAT1 in a multi-drug resistant *T. b. brucei* clone, B48, showed that this gene encoded a P1-type broad specificity purine nucleoside transporter with no ability to transport diminazene [31], and genomic analysis has shown that there is no direct orthologue of the P2/TbAT1 transporter in the genome of *T. congolense*. Regrettably, this has left us still without any insights into the urgent problem of DA resistance in animal trypanosomiasis. One urgent question beyond that of the resistance mechanism is that of cross-resistance with isometamidium, which has been reported repeatedly from the field [32,33], but inconsistently, as others report no cross-resistance and it used to be a rare occurrence [1,34,35]. Clearly, it is possible that trypanosomes may have developed resistance to both drugs independently over the decades and are now resistant to both by separate mechanisms. Thus, it is currently unclear whether other diamidine drugs should be considered for novel drug development for AAT because of the risk of cross-resistance with DA. This adds to the many reasons why it is important to understand the mechanism of DA resistance in animal-infective trypanosomes, alongside the need to identify screening tools for AAT drug resistance to assess the real levels of resistance in the field.

Here, several of these issues are addressed in an attempt to establish a hypothesis for DA resistance in *T. congolense*, and we report that DA resistance is not linked to reduced drug accumulation in *T. congolense*, nor necessarily to cross-resistance with isometamidium. However, we present evidence of stable mitochondrial changes as the basis for DA resistance. In sharp contrast to *T. brucei* spp, uptake of [^3^H]-DA by *T. congolense* was low affinity and inefficient and involves, at least in part, folic acid transporters.

## Results and Discussion

### *In vitro* induction of diminazene aceturate resistance in *T. congolense*

Four cultures of *T. congolense* TcoIL3000 were grown and passaged in parallel; a control without added drug and three separate, independent cultures in the presence of DA for adaptation to increasing concentrations, from which six clonal lines were obtained. In this study we used one clone from original line 4 (4C2), and two clones each from lines 5 (5C1, 5C2) and 6 (6C1, 6C3). Resistance was very slow to develop, as documented in S1 Fig. At the end of the adaptation, after 227 passages, the control culture tolerated 50 nM DA, whereas all the adapted cultures tolerated a maximum level of 800 nM DA, in what appeared to be at least two steps, with an initial plateau at 550 nM, where the strains were assessed for their level of DA resistance, and cross-resistance pattern.

At the point of adaptation to 550 nM, the level of DA resistance and the extent of any cross-resistance with the diamidine drugs pentamidine and DB829, and with oxaboroles AN7973 and SCYX7158, was tested using an assay based on the viability dye Alamar blue (resazurin), in which live trypanosomes (but not dead ones) reduce blue and non-fluorescent resazurin to pink, fluorescent resorufin. This assay yields a fluorescent output that is proportional to the number of cells in the well [36]. The results, summarized in Fig 1A, show that all the DA-Res strains display highly similar drug sensitivity patterns. Resistance to DA, as measured in this assay, was ~2.3-3-fold, and highly significant (Fig 1A for fold differences and P values, Table 1 for EC_50_ values). No cross-resistance with pentamidine was observed, but the level of resistance to DB829, a structurally closer analogue of DA, mirrored the resistance profile of DA very closely, demonstrating that cross-resistance to diamidines is not inevitable, but observed only with close structural analogues. Importantly, there was no cross-resistance with isometamidium, nor with either of the oxaboroles tested, AN7973 and SCYX7158. To further investigate the limits of cross-resistance with other diamidines, a series of bis-benzofuran analogues [37] were tested (Table 1). These analogues were chosen to explore the effect of variable linker length between the benzofuramidine end groups, varying inter-amidine distance as well as the flexibility of the molecule. All the benzofuramidines displayed low micromolar activity (Table 1) and there was no crossresistance with DA (Fig 1B). These results clearly show that the cross-resistance phenotype is very narrow and that many diamidines are still potentially useful for the treatment of DA-resistant AAT.

**Fig 1.**
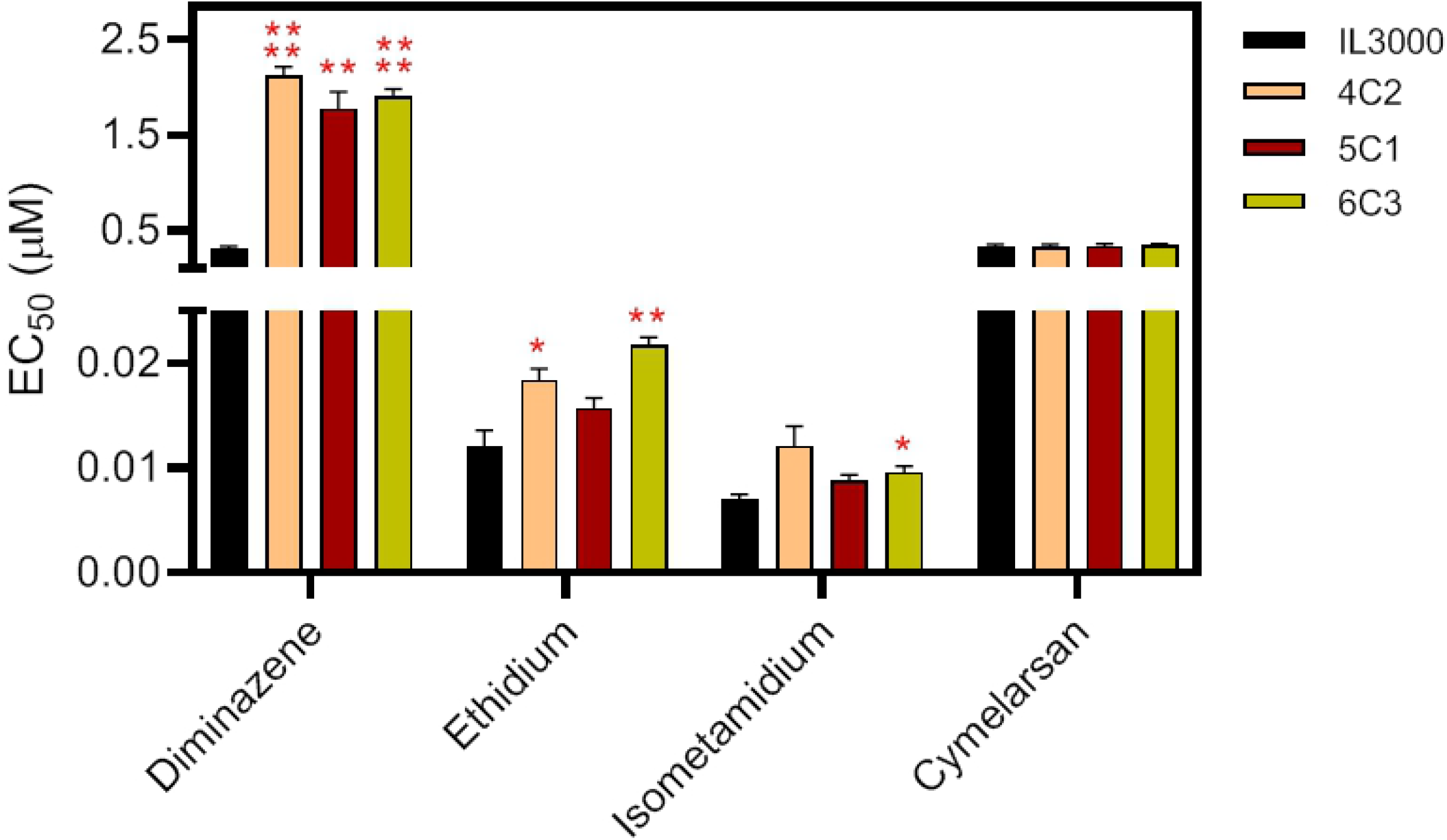
*In vitro* cross-resistance pattern of clones from three independent diminazene-adapted cultures with EC_50_ values given relative to that for the control strain, IL3000, adapted to growth in 550 nM DA. A. Cross-resistance to DB829 but not pentamidine, isometamidium, AN7973 or SCYX7158 was observed (n = 5 – 9). B. Cross-resistance to a series of bis-benzofuramidines was investigated – none of the strains were more than marginally resistant or sensitized to these compounds and the average EC_50_ of the 5 resistant strains was in all cases very close to the value for the IL3000 control. All structures are displayed in Table 1. Statistical significance between EC_50_s of a resistant line and the IL3000 control was determined using Student’s unpaired, two-tailed t-test; *, P<0.05; **, P<0.01; ***, P<0.001; ****, P<0.0001.

**Table 1.**
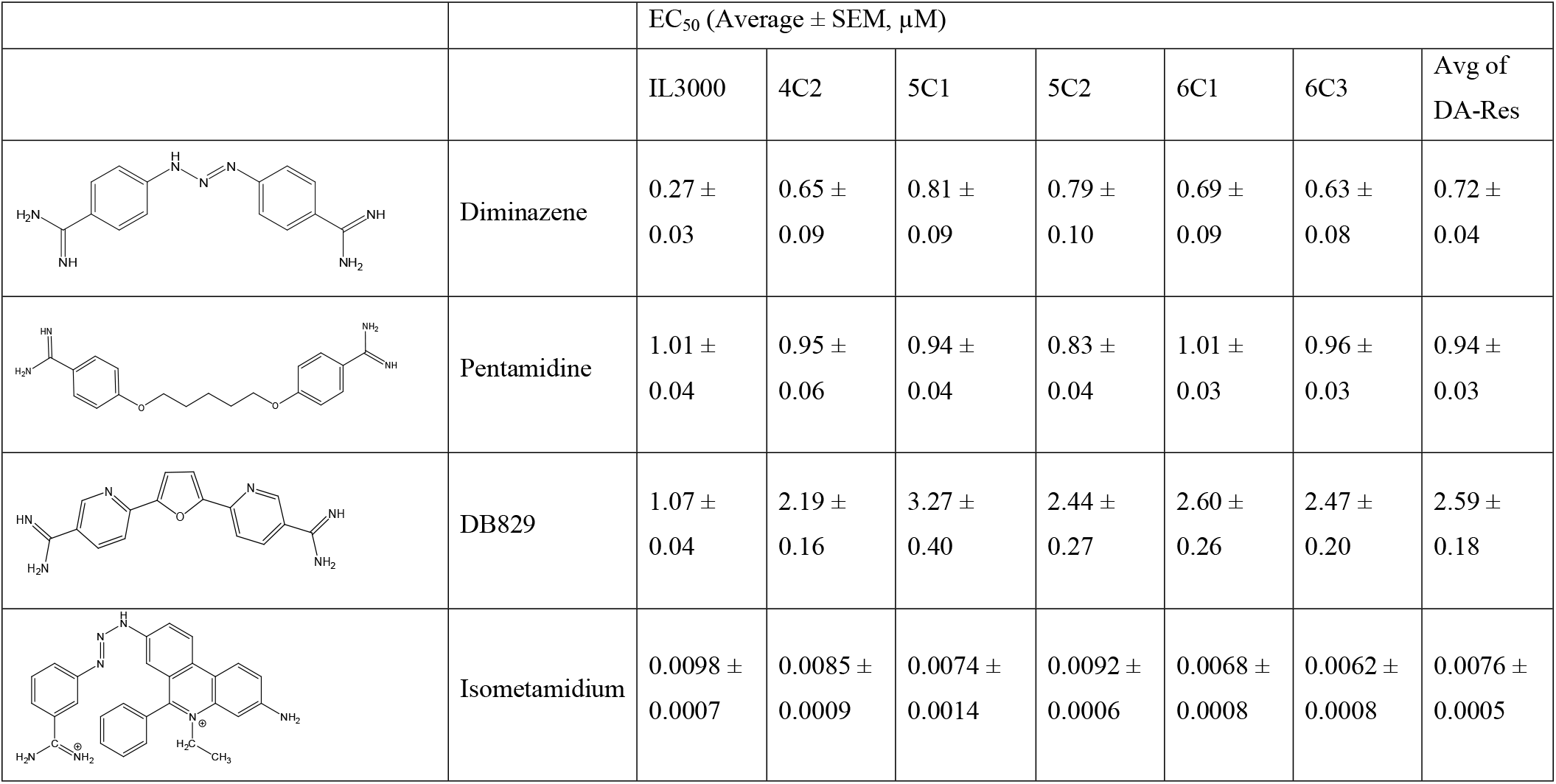
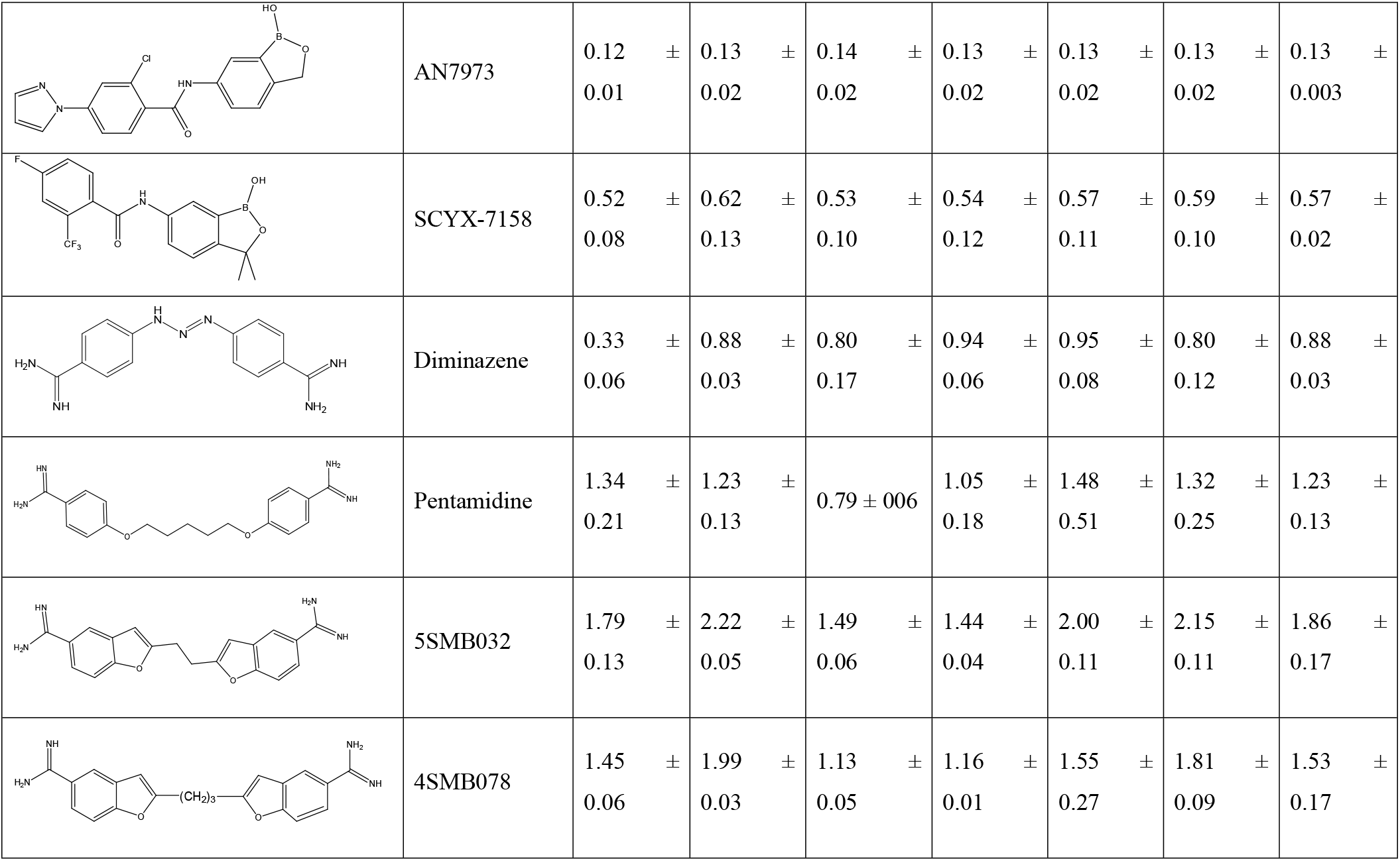
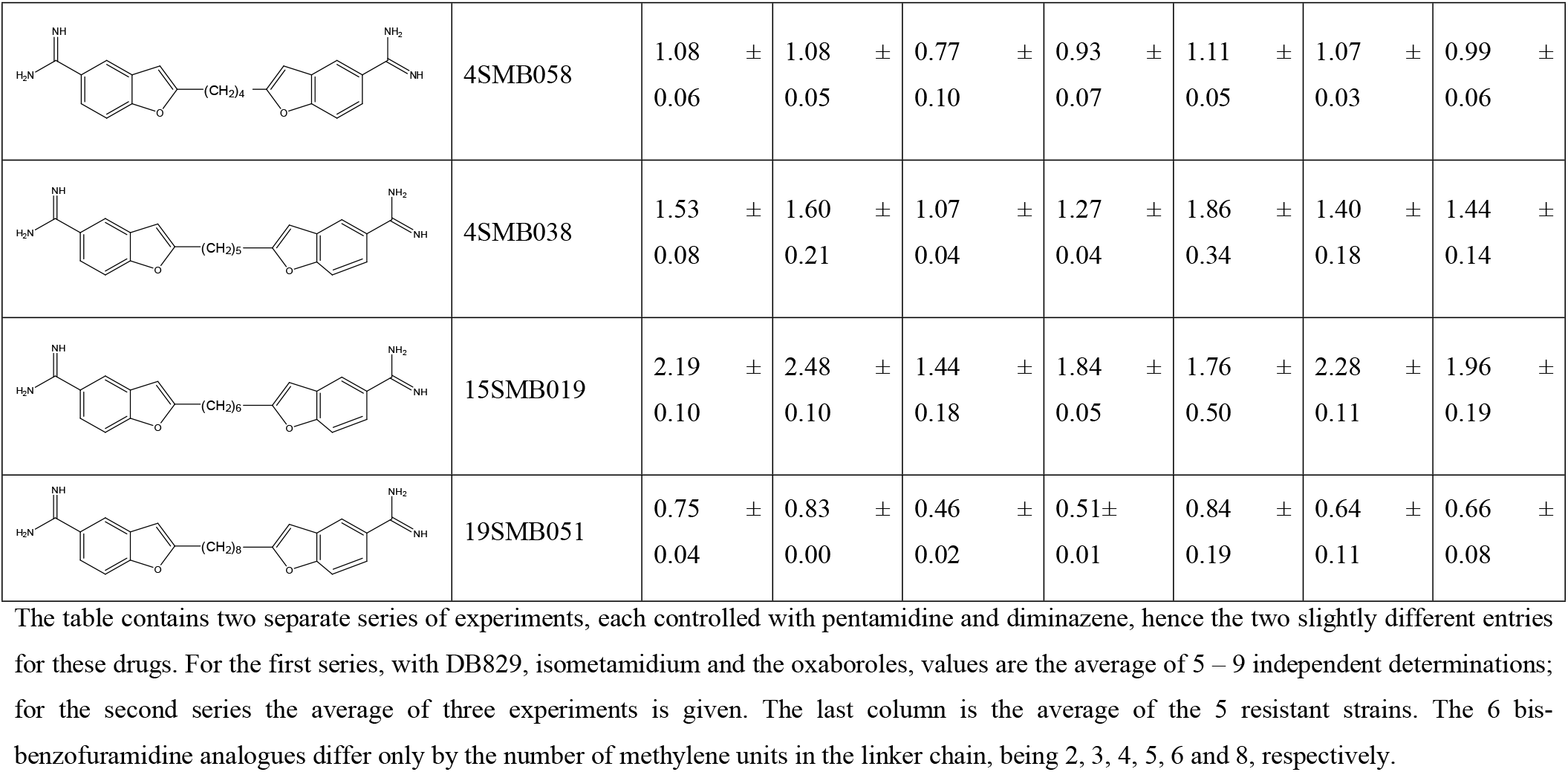
*In vitro* cross-resistance pattern for diminazene-adapted *T. congolense* strains and the parental strain IL3000.

|  |  | EC<sub>50</sub> (Average ± SEM, µM) |  |  |  |  |  |  |
| --- | --- | --- | --- | --- | --- | --- | --- | --- |
|  |  | IL3000 | 4C2 | 5C1 | 5C2 | 6C1 | 6C3 | Avg of DA-Res |
|  | Diminazene | 0.27 ± 0.03 | 0.65 ± 0.09 | 0.81 ± 0.09 | 0.79 ± 0.10 | 0.69 ± 0.09 | 0.63 ± 0.08 | 0.72 ± 0.04 |
|  | Pentamidine | 1.01 ± 0.04 | 0.95 ± 0.06 | 0.94 ± 0.04 | 0.83 ± 0.04 | 1.01 ± 0.03 | 0.96 ± 0.03 | 0.94 ± 0.03 |
|  | DB829 | 1.07 ± 0.04 | 2.19 ± 0.16 | 3.27 ± 0.40 | 2.44 ± 0.27 | 2.60 ± 0.26 | 2.47 ± 0.20 | 2.59 ± 0.18 |
|  | Isometamidium | 0.0098 ± 0.0007 | 0.0085 ± 0.0009 | 0.0074 ± 0.0014 | 0.0092 ± 0.0006 | 0.0068 ± 0.0008 | 0.0062 ± 0.0008 | 0.0076 ± 0.0005 |
|  | AN7973 | 0.12 ± 0.01 | 0.13 ± 0.02 | 0.14 ± 0.02 | 0.13 ± 0.02 | 0.13 ± 0.02 | 0.13 ± 0.02 | 0.13 ± 0.003 |
|  | SCYX-7158 | 0.52 ± 0.08 | 0.62 ± 0.13 | 0.53 ± 0.10 | 0.54 ± 0.12 | 0.57 ± 0.11 | 0.59 ± 0.10 | 0.57 ± 0.02 |
|  | Diminazene | 0.33 ± 0.06 | 0.88 ± 0.03 | 0.80 ± 0.17 | 0.94 ± 0.06 | 0.95 ± 0.08 | 0.80 ± 0.12 | 0.88 ± 0.03 |
|  | Pentamidine | 1.34 ± 0.21 | 1.23 ± 0.13 | 0.79 ± 0.06 | 1.05 ± 0.18 | 1.48 ± 0.51 | 1.32 ± 0.25 | 1.23 ± 0.13 |
|  | 5SMB032 | 1.79 ± 0.13 | 2.22 ± 0.05 | 1.49 ± 0.06 | 1.44 ± 0.04 | 2.00 ± 0.11 | 2.15 ± 0.11 | 1.86 ± 0.17 |
|  | 4SMB078 | 1.45 ± 0.06 | 1.99 ± 0.03 | 1.13 ± 0.05 | 1.16 ± 0.01 | 1.55 ± 0.27 | 1.81 ± 0.09 | 1.53 ± 0.17 |
|  | 4SMB058 | 1.08 ± 0.06 | 1.08 ± 0.05 | 0.77 ± 0.10 | 0.93 ± 0.07 | 1.11 ± 0.05 | 1.07 ± 0.03 | 0.99 ± 0.06 |
|  | 4SMB038 | 1.53 ± 0.08 | 1.60 ± 0.21 | 1.07 ± 0.04 | 1.27 ± 0.04 | 1.86 ± 0.34 | 1.40 ± 0.18 | 1.44 ± 0.14 |
|  | 15SMB019 | 2.19 ± 0.10 | 2.48 ± 0.10 | 1.44 ± 0.18 | 1.84 ± 0.05 | 1.76 ± 0.50 | 2.28 ± 0.11 | 1.96 ± 0.19 |
|  | 19SMB051 | 0.75 ± 0.04 | 0.83 ± 0.00 | 0.46 ± 0.02 | 0.51 ± 0.01 | 0.84 ± 0.19 | 0.64 ± 0.11 | 0.66 ± 0.08 |

The table contains two separate series of experiments, each controlled with pentamidine and diminazene, hence the two slightly different entries for these drugs. For the first series, with DB829, isometamidium and the oxaboroles, values are the average of 5 – 9 independent determinations; for the second series the average of three experiments is given. The last column is the average of the 5 resistant strains. The 6 bis-benzofuramidine analogues differ only by the number of methylene units in the linker chain, being 2, 3, 4, 5, 6 and 8, respectively.

The DA-resistance of the five DA-resistant clones was then increased via further *in vitro* exposure until they could tolerate growth at 800 nM DA. The *in vitro* growth rates of these strains consistently appeared to be slightly slower than that of the drug-sensitive parental strain (Fig 2), although this reached statistical significance only for the 48 h and 64 h points of clones 5C2 and 6C1 (P<0.05, unpaired student’s t-test). The EC_50_ values, at ~2.2 μM, were similar for all clones but significantly higher than when they were measured before the cloning, at 550 nM DA medium concentration. This level of resistance, approximately 9-fold, was stable for at least 3 months of *in vitro* culturing in the absence of drug pressure (Fig 3). Cross-resistance to DB829 was again observed in all clones (~5-fold), whereas sensitivity to pentamidine was identical in all clones including wildtype IL3000. Interestingly, sensitivity to AN7973 was slightly but significantly higher in most DA-Res clones (Fig 3). Crossresistance with phenanthridines isometamidium and ethidium bromide, and with the melaminophenyl arsenical cymelarsan was also tested for selected clonal lines. A slight loss of sensitivity to phenanthridines was consistently observed and reached significant (but <2-fold) levels in some clones. No change in cymelarsan sensitivity was observed (Fig 4).

**Fig 2.**
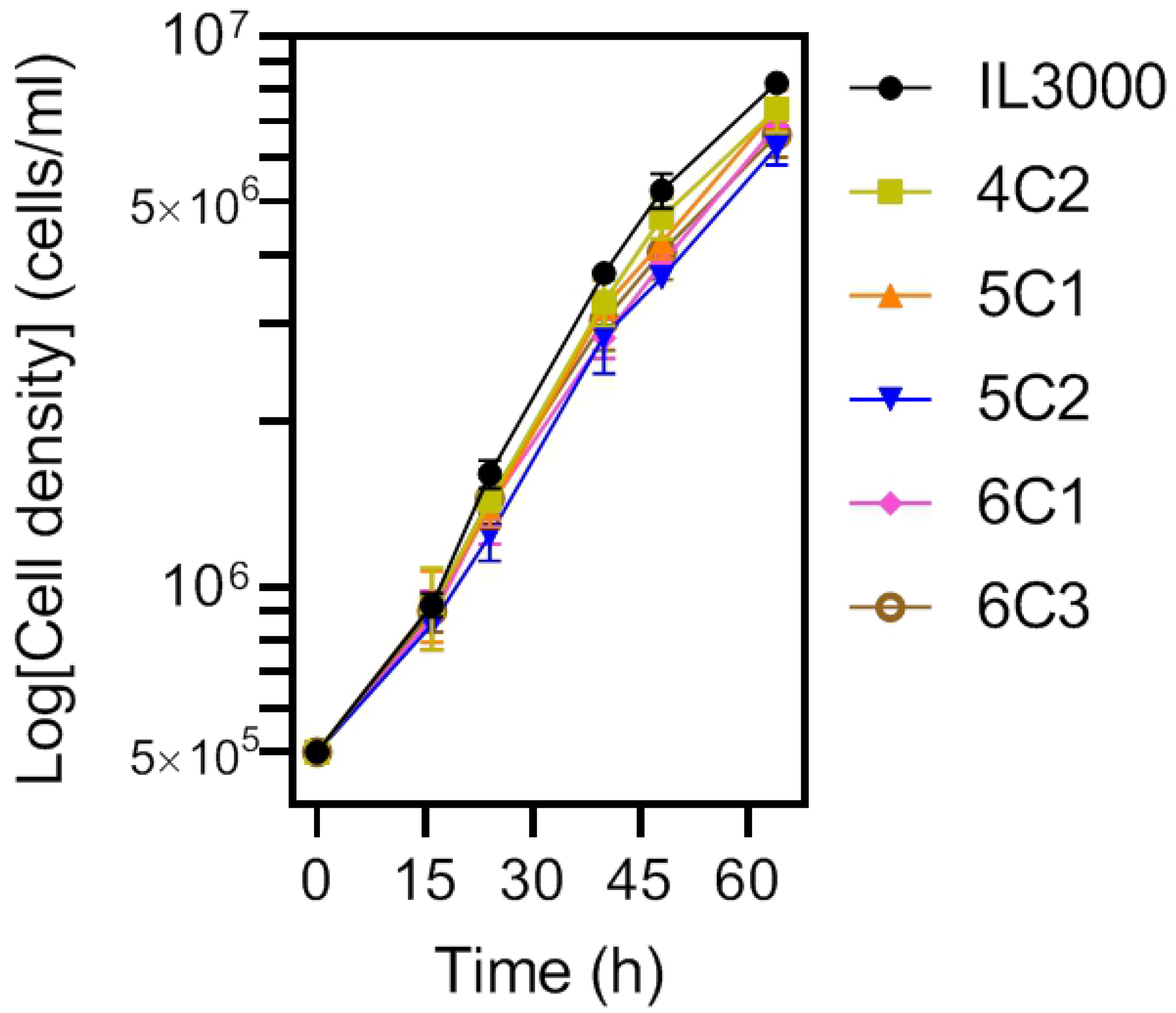
Growth curve of diminazene-sensitive TcoIL3000 and 5 different diminazene aceturate-resistant *T. congolense* clones. The cells were cultured at 34 °C and 5% CO_2_ in TcBSF-3 medium for up to 64 h in a 6-well plate and counted periodically using a haemocytometer. The graph shows the average and SEM of three independent repeats performed with different cultures. When the error bars are not shown they fall inside the symbol.

**Fig 3.**
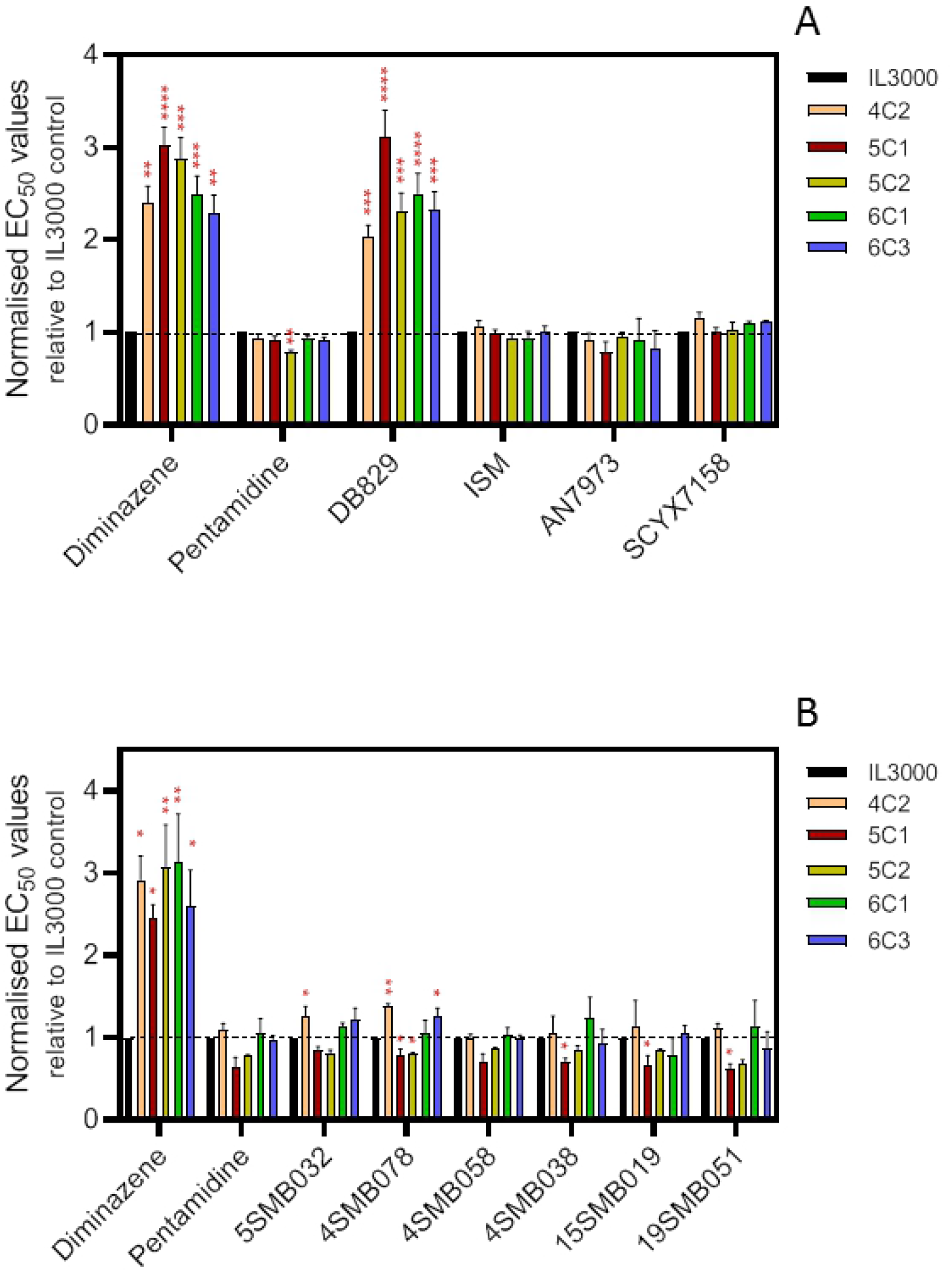
Stability of resistance phenotype in diminazene-adapted *T. congolense* clones. EC_50_ values were obtained using the Alamar blue assay. All bars represent the average and SEM of 4 independent determinations. Statistical significance between EC_50_s of a resistant line and the IL3000 control was determined using Student’s unpaired, two-tailed t-test; *, P<0.05; **, P<0.01; ***, P<0.001; ****, P<0.0001. IL3000 = wild-type control; A = grown in the presence of 800 nM diminazene; B = passaged for 3 months in the absence of diminazene drug pressure but previously grown in 800 nM DA.

**Fig 4.**
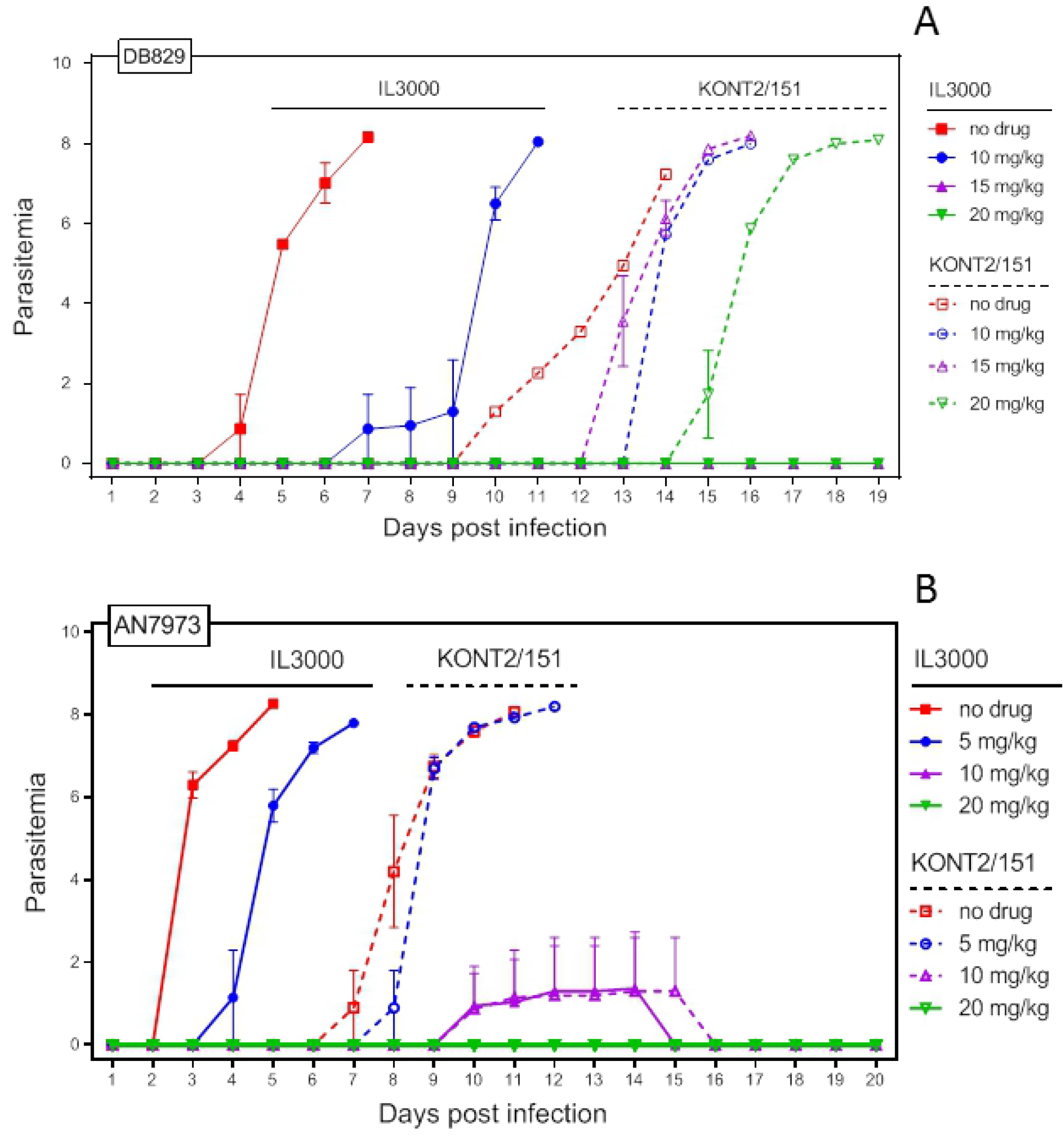
Alamar blue assay with IL3000 and three DA-Res clones from culture, grown in the absence of drug pressure. All bars represent the average and SEM of 3 independent determinations. Statistical significance between EC_50_s of a resistant line and the IL3000 control was determined using Student’s unpaired, two-tailed t-test; *, P<0.05; **, P<0.01; ***, P<0.001; ****, P<0.0001.

### *In vivo* confirmation of cross-resistance profile

In order to confirm that the *in vitro* pattern of cross-resistance held up for *in* vivo infections, including for a field-derived DA-resistant strain, we infected groups of 6 mice that were treated i.p. with either DB829 (Fig 5A) or AN7973 (Fig 5B), using single doses of 5, 10 or 20 mg/kg bw (two separate independent experiments with each drug, each experiment n=6). For the resistant filed isolate we used stain KONT2/151 [38]. IL3000 was more virulent than KONT2/151, as is often observed with cultured trypanosome strains, and was sensitive to both DB829 and AN7973. DB829 delayed parasitemia and death with a dose of 10 mg/kg and was curative at 15 and 20 mg/kg, whereas AN7973 delayed parasitemia at 5 mg/kg and cured with 10 or 20 mg/kg. The DA-resistant field isolate was equally sensitive to the oxaborole but resistant to DB829, as even the highest dose, 20 mg/kg, merely delayed the onset of parasitemia.

**Fig 5.**
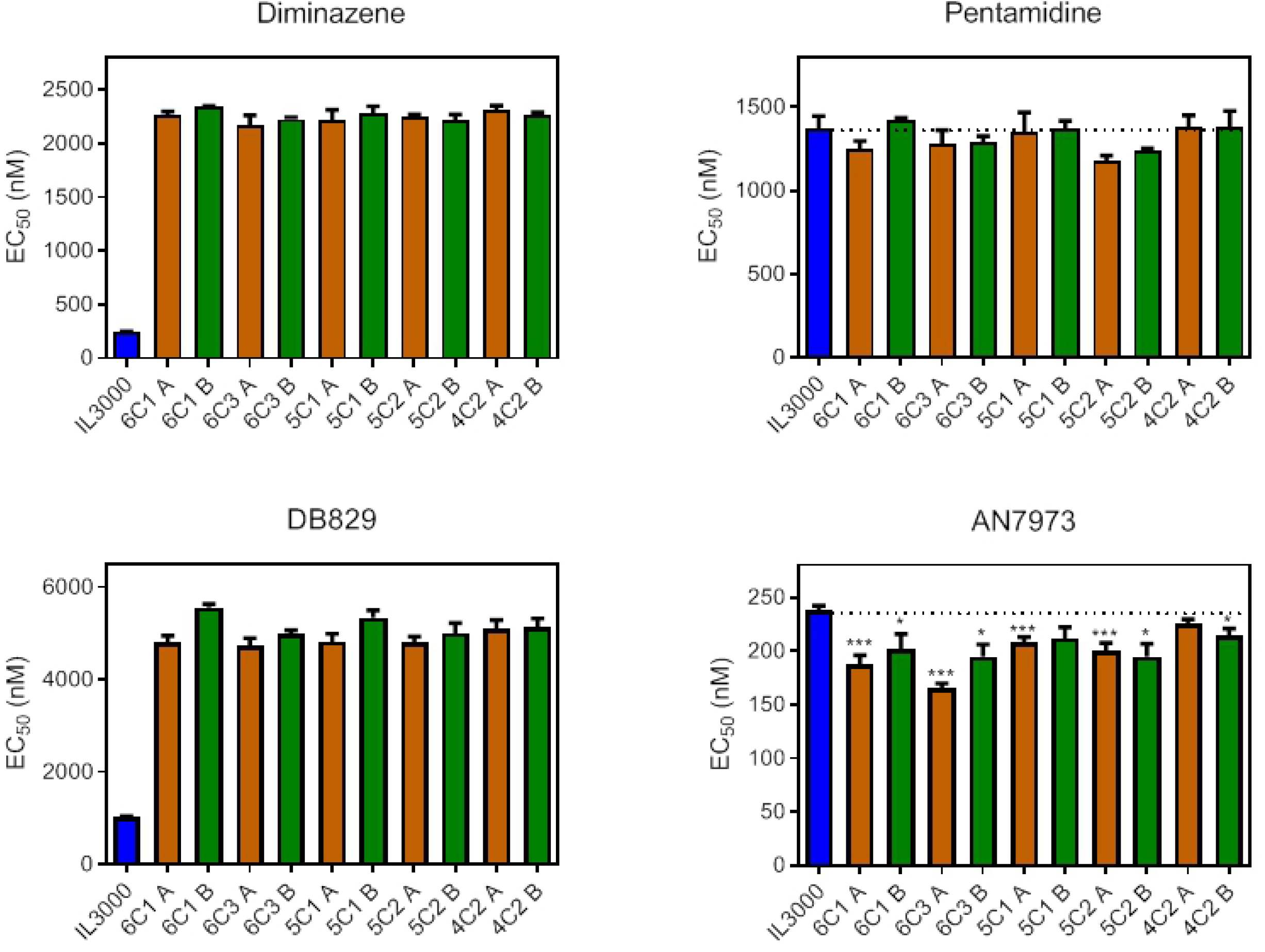
*In vivo* analysis of drug sensitivity for drug sensitive *T. congolense* clone IL3000 and diminazene resistant isolate *T. congolense* KONT2/151. A. Infected mice were treated with the indicated doses of DB829 or with vehicle (no-drug controls). Each group consisted of six mice. Parasitaemia was determined daily from blood taken from tail-pricks and shown as average ± SEM. Mice infected with IL3000 are represented by solid symbols and lines; KONT2/151 by open symbols and dotted lines. (B) Like A, but the treatment was with AN79373. The results shown are single experiments but representative of 2 very similar experiments with essentially the same outcomes.

### Screen for potential diminazene transport inhibitors by DB75 fluorescence microscopy

In *T. b. brucei*, fluorescent diamidines such as DB75, a close structural analogue of DA and DB829, have previously been used to monitor uptake and cellular distribution of such compounds [22]. In those experiments, the rate at which the kinetoplast (first) and the nucleus (second) became visible in the blue wavelengths (λ=405/435 nm for excitation and emission, respectively) was much delayed in resistant strains, and in the presence of transport inhibitors [22]. Here, the technique was used to screen for potential transport substrates that competitively inhibit the uptake of the DB75 in animal trypanosome species. The utility of the technique was first confirmed in *ex vivo T. b. brucei* in mouse blood. Murine blood cells did not appear to significantly take up 10 μM DB75, ensuring a low background, but in *T. b. brucei*, parasite outlines would become visible after 4 min, followed by kinetoplasts at 5-6 min and finally nuclei were faintly stained by 8 min and bright by 15 min. These stages were all observed and scored in the presence of potential inhibitors. As expected, based on current *T. brucei* models [25,26], the polyamines putrescine, spermine and spermidine (500 μM) had no discernible effect on DB75 staining, although the polyamine transporters of *Leishmania* spp and *T. cruzi* have been implicated in possible pentamidine uptake [39–41]. In contrast, DA and the shorter pentamidine analogue propamidine both dose-dependently delayed DB75 fluorescence (S1 Table); both are known substrates of the TbAT1/P2 aminopurine transporter [7,42] that is the sole uptake mechanism for DB75 in *T. brucei* [43].

Because of the involvement of the TbAT1/P2 purine transporter in DA uptake by *T. b. brucei*, we tested a range of purine and pyrimidine nucleosides and nucleobases for delaying DB75 cell entry in *T. congolense*. Adenine and adenosine did not appear to slow DB75 fluorescence up to 1 mM but were toxic to the cells at concentrations of 200 μM and above. Other purines (inosine, guanosine, hypoxanthine, guanine, xanthine) or pyrimidines (uridine, cytidine, thymidine, uracil, cytosine) likewise failed to inhibit DB75 uptake; moreover, polyamines and amino acids (500 μM) were similarly ineffective in this assay (S1 Table). Glycerol was also tested at up to 20 mM because some diamidines including pentamidine are taken up by an aquaglyceroporin in *T. brucei* [9,27], but the data indicated that glycerol also does not compete for the *T. congolense* DB75 transporter. However, the diamidines DA and, particularly, propamidine and pentamidine, dose-dependently (100 μM – 500 μM) delayed DB75 fluorescence, and so did folic acid, from 200 μM to 500 μM (Fig 6). *T. vivax* was also assayed by this method and similar results were found, with no effect from a series of purines and pyrimidines, glycerol, and the cationic amino acids lysine and arginine, but dosedependent inhibition by pentamidine and DA. For this parasite, there appeared to be some minor fluorescence delay in the presence of 500 μM folic acid or biopterin, but not at lower concentrations, and possibly in the presence of the polyamines spermidine and putrescine (S1 Table).

**Fig 6.**
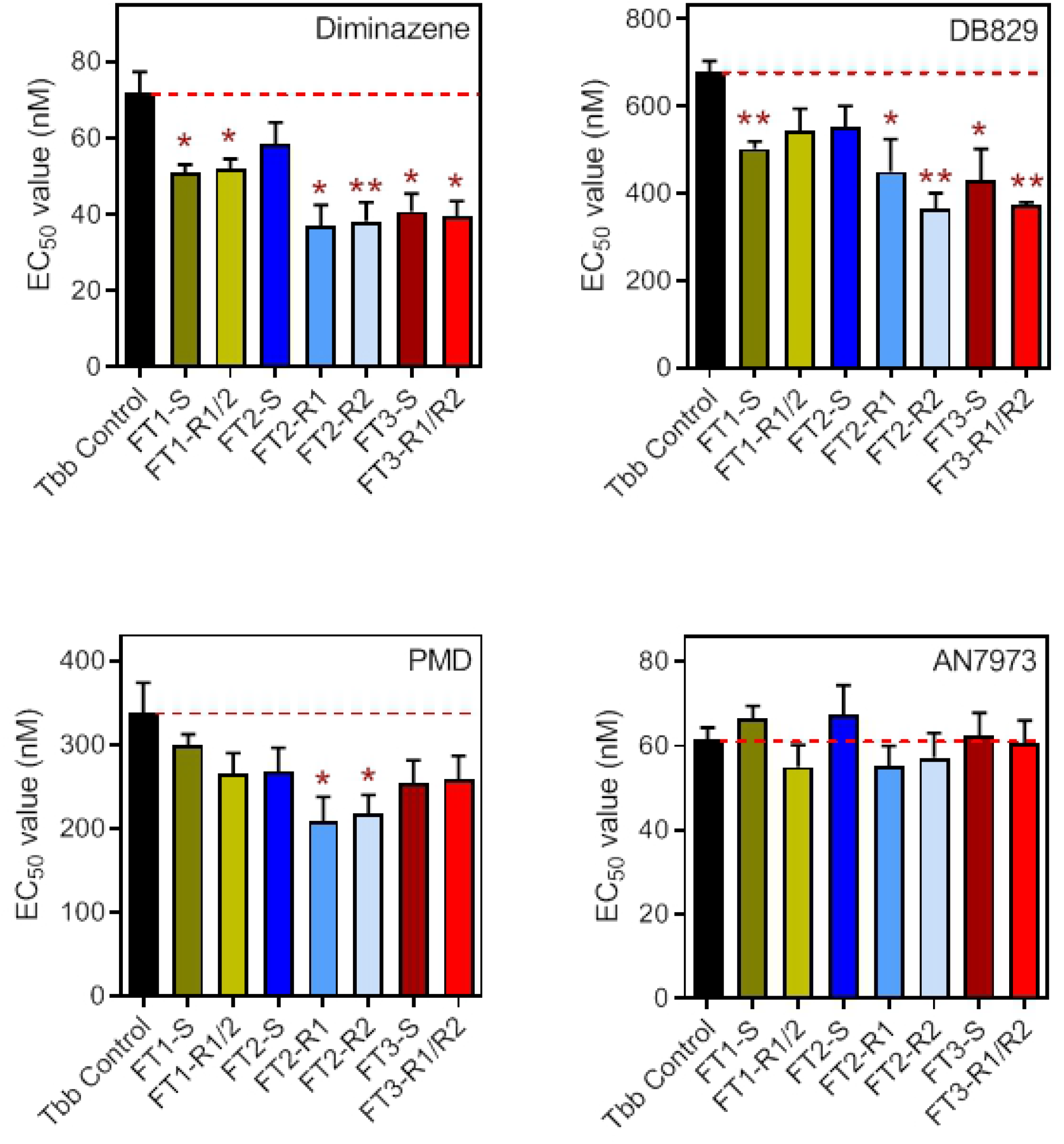
DB75 fluorescence in *ex vivo T. congolense* trypomastigotes. Parasites were obtained from infected mice and incubated with 10 μM DB75. Fluorescence was monitored under an Axiosope fluorescence microscope (Zeiss) at an excitation wavelength of 330 nm and an emission wavelength of 400 nm. Images were obtained with Openlab imaging software (Improvision, Coventry, UK). The brightfield and fluorescence images shown were obtained after 12 min incubation, with the nucleus (N) and kinetoplast (K) of two parasites in the lefthand panel visible as blue spots (no inhibitor control) and no fluorescence in the right-hand panel (500 μM DA). These images are representative of similar images taken in approximately 2 dozen such experiments.

### Uptake of [^3^H]-Diminazene by DA-sensitive and -resistant *T. congolense* clones

In *T. brucei*, resistance to DA is linked to loss of the TbAT1/P2 transporter [6,11,25]. While *T. congolense* does not have an equivalent transporter [44], drug resistance in trypanosomatids has very often been associated with the loss of transporters at the plasma membrane [10,12,45] and we thus investigated whether uptake of [^3^H]-DA was altered in the resistant clones.

Initial experiments with various concentrations of [^3^H]-DA, trying to standardize linear uptake conditions, were unsuccessful. Uptake was very slow, the label displayed high background from binding to the outside of the cell and a true linear phase of uptake could not be established with the requisite level of confidence and reproducibility. Therefore, accumulation of 0.1 μM [^3^H]-DA was monitored over 30 minutes in IL3000 and two of the derived DA-resistant clones, 4C2 and 6C3. This showed that even over half an hour there was no clear difference in the rate of accumulation of [^3^H]-DA in any of the three cell lines and that resistance is therefore very unlikely to be the result of induced changes in drug uptake rate (Fig 7A). Nor was there significant efflux from the cells, after preincubation with 3H]-DA for 30 min followed by incubation of up to 30 minutes in fresh medium without label or diminazene. Although the slope of both the IL3000 WT and the 4C2 clone trended downwards, in each of 4 such experiments the slopes were not significantly different from zero, nor different from each other (F-test, P>0.05). Fig 7B shows the average of 4 experiments, which still shows a non-significant downward trend and no difference between the slope of the sensitive and resistant lines. In order to better understand why DA-resistance in *T. congolense*, in contrast to *T. b. brucei*, is not associated with loss of import or gain of efflux function, the mechanism by which DA is internalized by *T. congolense* was further investigated.

**Fig 7.**
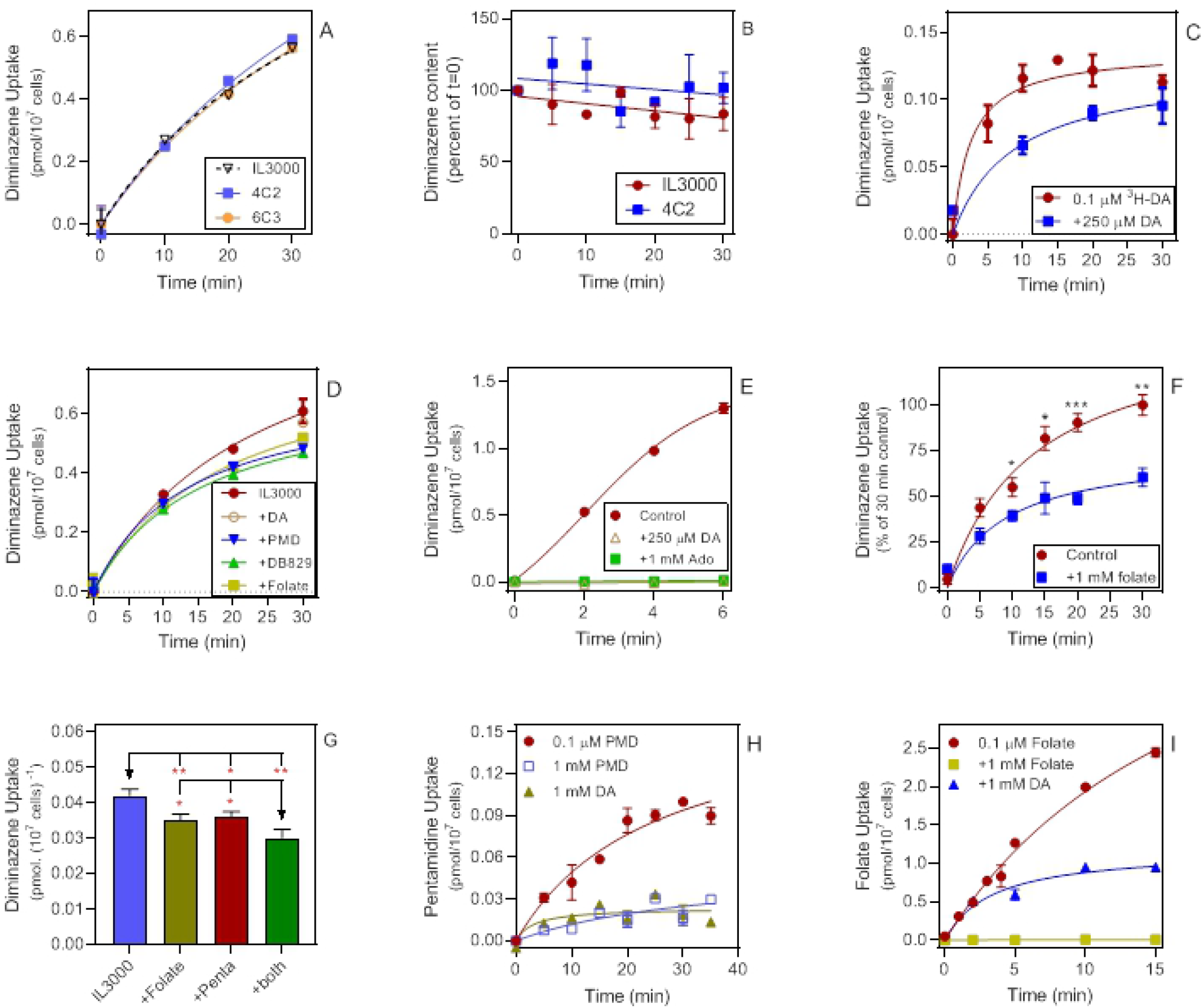
Investigation into the uptake of diminazene by *T. congolense*. A. Uptake of 0.1 μM [^3^H]-DA by IL3000 and two derived DA-adapted clones. Uptake of DA was monitored for 30 minutes in *in vitro* grown cells of *T. congolense* IL3000, 4C2 and 6C3. There were no statistically significant differences between the three strains at any of the timepoints. B. Efflux of [^3^H]-DA from *T. congolense* IL3000 and from clone 4C2, after loading with 0.1 μM [^3^H]-DA for 30m minutes, followed by a wash into fresh assay buffer. Samples were taken at the indicated time points after resuspension into fresh buffer. The data presented is the average of each time point of 4 independent experiments, each performed in triplicate. The two slopes, calculated by linear regression, were not significantly different from each other (P=0.84, F-test), nor significantly different from zero (P=0.14 and 0.46 for IL3000 and 4C2, respectively). C. IL3000 parasites were isolated from mice and incubated with 0.1 μM [^3^H]-DA with and without 250 μM unlabelled DA. D. IL3000 parasites from culture were incubated with 0.1 μM [^3^H]-DA label in the presence or absence of 100 μM DA, 100 μM pentamidine (PMD), 100 μM DB829 or 1 mM folic acid. E. Uptake of 0.1 μM [^3^H]-DA by bloodstream forms of *T. b. brucei* clone B48 transfected with *TbAT1* without inhibitor or in the presence of 250 μM DA or 1 mM adenosine (Ado). F. Uptake of 0.1 μM [^3^H]-DA by *T. congolense* IL3000 from culture, in the presence and absence of 1 mM folic acid, expressed as % of uptake in the absence of folic acid at 30 min. The average and SEM of 6 independent experiments, each performed in triplicate, is shown. G. *T. congolense* IL3000 from culture were incubated with 0.1 μM [^3^H]-DA for 20 min with either no inhibitor, 1 mM folic acid, 100 μM pentamidine, or both together. The level of inhibition was virtually identical with either of the two inhibitors, but significantly higher (P<0.05) when the medium contained both. Bars represent the average and SEM of a triplicate determination. H. Uptake of 0.1 μM [^3^H]-pentamidine by *T. congolense* IL3000 from culture in the presence or absence of 1 mM pentamidine (PMD) or diminazene aceturate (DA). One representative experiment in triplicate is shown. I. Uptake of 0.1 μM [^3^H]-folic acid by *T. congolense* IL3000 from culture, and the effects of 1 mM folic acid or 1 mM DA. One representative experiment in triplicate is shown. All experiments were performed in triplicate and the average and SEM are shown. When error bars are not shown, they fall inside the symbol. Statistical significance was calculated using Student’s unpaired t-test: *, P<0.05; **, P<0.01; ***, P<0.001).

Uptake of 0.1 μM [^3^H]-DA was poorly saturable by even a large excess of unlabeled DA (Fig 7C), indicating that DA enters *T. congolense* by a low affinity and/or non-saturable process. Indeed, out of several potential inhibitors or substrates of the DA uptake, the process was consistently (but always partially) inhibited by 100 μM pentamidine, 1 mM folate and 100 μM DB829 (Fig 7D). This is in sharp contrast to DA uptake in *T. brucei* which is completely saturable and 100% inhibited by a competitive substrate of the TbAT1 transporter (Fig 7E). The inhibition with folate was confirmed and quantified in a series of experiments with added time points, and found to be highly significant (Fig 7F). Moreover, in a separate experiment it was tested whether the inhibition by folate and pentamidine was additive and the data indicated that this was the case (Fig 7G), suggesting that in *T. congolense* DA is taken up by at least two separate low affinity transporters, one sensitive to folate and another that is sensitive to pentamidine. However, it must be noted that the inhibition by both compounds was modest in this experiment and that the joint administration still did not inhibit [^3^H]-DA uptake by even 50%. The complete lack of cross-resistance with pentamidine, despite high concentrations of pentamidine partially inhibiting DA uptake, is consistent with DA resistance not being the result of changes in its uptake rate. Indeed, the overlap in transport mechanisms of DA and pentamidine was even more clear from the inhibition of [^3^H]-pentamidine uptake by DA in wild-type IL3000 (Fig 7H). To further investigate whether (a) folate transporter(s) might contribute to DA uptake in *T. congolense*, we measured the uptake of [^3^H]-folate and found this to be partially inhibited by 1 mM DA (Fig 7I).

#### *T. congolense* folate transporters are engaged in diminazene uptake

Three genes encoding putative folate transporters of *T. congolense* were identified via homology searches in TriTrypDB in the original *T. congolense* IL3000 genome and amplified by PCR with Phusion proof-reading polymerase from IL3000 and from the DA-resistant isolates KONT2/133 and KONT2/151 (sequences, GeneIDs and GenBank accession numbers in Table 2); all three strains are Savannah-type *T. congolense*. The genes were cloned into pGEM-T Easy for amplification in *E. coli* and for each gene at least 4 independent clones were sequenced. The gene sequences from the two KONT strains, both from Cameroon, were almost identical to each other, but displayed some SNPs relative to IL3000 (Table 3). For folate transporter 1 (*TcoFT1*, 592 aa) there were two identical, conservative SNPs of IL3000 alanine to KONT serine (positions 49 and 331). For FT2 (640 aa) there were 8 shared non-synonymous SNPs in the two KONT strains and one unique one in each, and for FT3, both KONT strains carried the same 15 amino acid changes relative to IL3000 (S2 Fig); a table with the percentage identity and similarity between these genes is included in S2 Table. The genes were cloned into the pHD1336 expression vector and transfected into *T. b. brucei* strain B48. This clonal line lacks both the *TbAT1*/P2 transporter and a wild-type *TbAQP2* gene, and is thus highly resistant to DA, the furamidines, pentamidine and melaminophenyl arsenicals [8,27,43]. As the sensitivity to these drugs is completely restored upon re-introduction of the transporters [27,44] this is an ideal system to test potential diamidine transporters from other kinetoplastids. Clonal lines were obtained by limiting dilution; the correct integration of the expression constructs was verified by PCR.

**Table 2.**
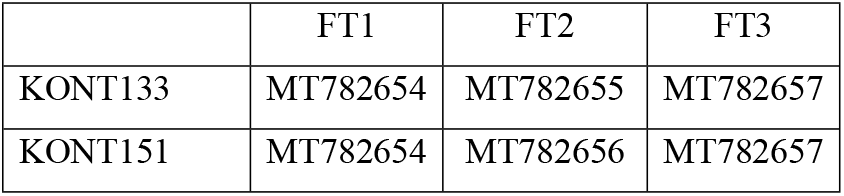
GenBank Accession Numbers of the *T. congolense* folate transporter genes.

|  | FT1 | FT2 | FT3 |
| --- | --- | --- | --- |
| KONT133 | MT782654 | MT782655 | MT782657 |
| KONT151 | MT782654 | MT782656 | MT782657 |

**Table 3.** Number of non-homologous SNPs in *T. congolense* folate transporter genes, relative to IL3000.

|  | FT1 | FT2 | FT3 |
| --- | --- | --- | --- |
| KONT133 | 2 | 9 | 15 |
| KONT151 | 2 | 8 | 15 |

Expression of the *T. congolense* folate transporters in *T. b. brucei* B48 confirmed that these transporters have some capacity to take up diamidines, but not the oxaborole AN7973 (Fig 8). Only FT2 of IL3000 did not significantly sensitize to any of the drugs tested, although its diamidine EC_50_ values also trended to be lower than the control (B48 transfected with the ‘empty vector’). Sensitization to DA and DB829 was almost identical, but only the FT2 transporters of the KONT strains significantly sensitized to pentamidine. In all cases the sensitization was modest, reaching ~50% with DA and DB829, and <40% for pentamidine, which is consistent with the transport experiments with TcIL3000, in that the transporters appear to display significant and yet very limited diamidine uptake capacity. For comparison, the expression of TbAT1 in B48 cells, using the same expression vector and assays, resulted in ~200-fold sensitization to DA and pentamidine instead of 2-fold [44]. The data presented in Fig 8 do not suggest that the two KONT strains are DA resistant because of changes in DA uptake – or at least not by their folate transporters. With respect to DA and DB829, the IL3000 and KONT FT1 and FT3 performed identically, but the KONT FT2 clearly sensitized stronger than the IL3000 FT2.

**Fig 8.**
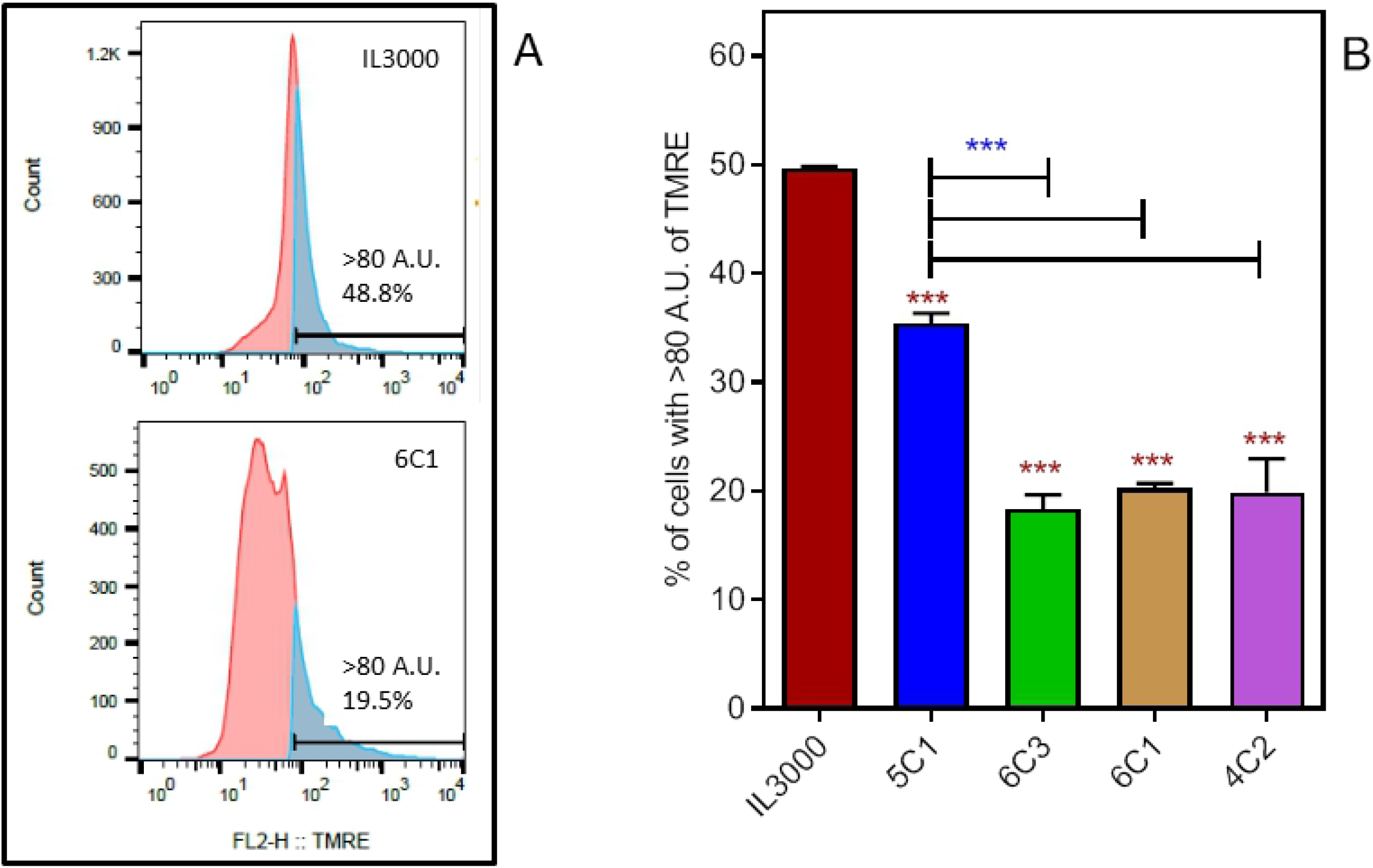
Drug sensitivity profile of *T. b. brucei* strain B48 (Tbb Control) and the same strain transfected with folate transporters 1-3 (FT1-3) of IL3000 (S), KONT2/133 (R1) or KONT2/151 (R2). EC_50_ values were determined using the Alamar blue assay. Bars represent the average and SEM of three independent experiments. Significant differences relative to the Tbb Control were calculated using Student’s unpaired t-test: *, P<0.05; **, P<0.01. PMD, pentamidine.

#### The mitochondrial membrane potential is diminished in DA-resistant *T. congolense* clones

As DA resistance is apparently not linked to changes in DA uptake, it was hypothesized that an intracellular rather than a cell surface change must be responsible for the resistance phenotype. Moreover, DA is a DNA minor grove binder and resistance arising due to mutations in a target protein are therefore not expected. Our observations with DB75 fluorescence microscopy showed that such drugs accumulate first in the kinetoplast, as reported for other species [22,46]. This requires rapid uptake into the mitochondrion, which is driven by the mitochondrial membrane potential Ψm as DA, DB75 and other diamidines are dications. We therefore investigated whether Ψm was altered in the resistant cell lines. Fluorescence microscopy after DAPI staining confirmed that the kinetoplast was present and did not appear altered or damaged in the DA-resistant clones (S3 Fig).

It is already well established that (di)cationic trypanocides that accumulate in the trypanosome’s mitochondrion cause a reduction in Ψm, both through the very fact that they are cations, and through disruption of mitochondrial processes (e.g. [23,24,47,49]). However, the more important question here was whether, as (part of) the adaptation to DA, the cells had permanently lowered their Ψm, as previously reported for isometamidium [20,21]. The DARes parasites did present flow cytometry profiles that had shifted significantly to lower fluorescence and were broader than the control IL3000 cells (Fig 9A). For quantification, the peak of the IL3000 profile was set at 80 Arbitrary Units (AU). Taking the percentage of cells with fluorescence >80 AU as a measure for Ψm [47,48], a highly significant decrease was found in all four DA-Res cell lines tested, although the extent of the Ψm decrease was not identical in all cell lines (Fig 9B).

**Fig 9.**
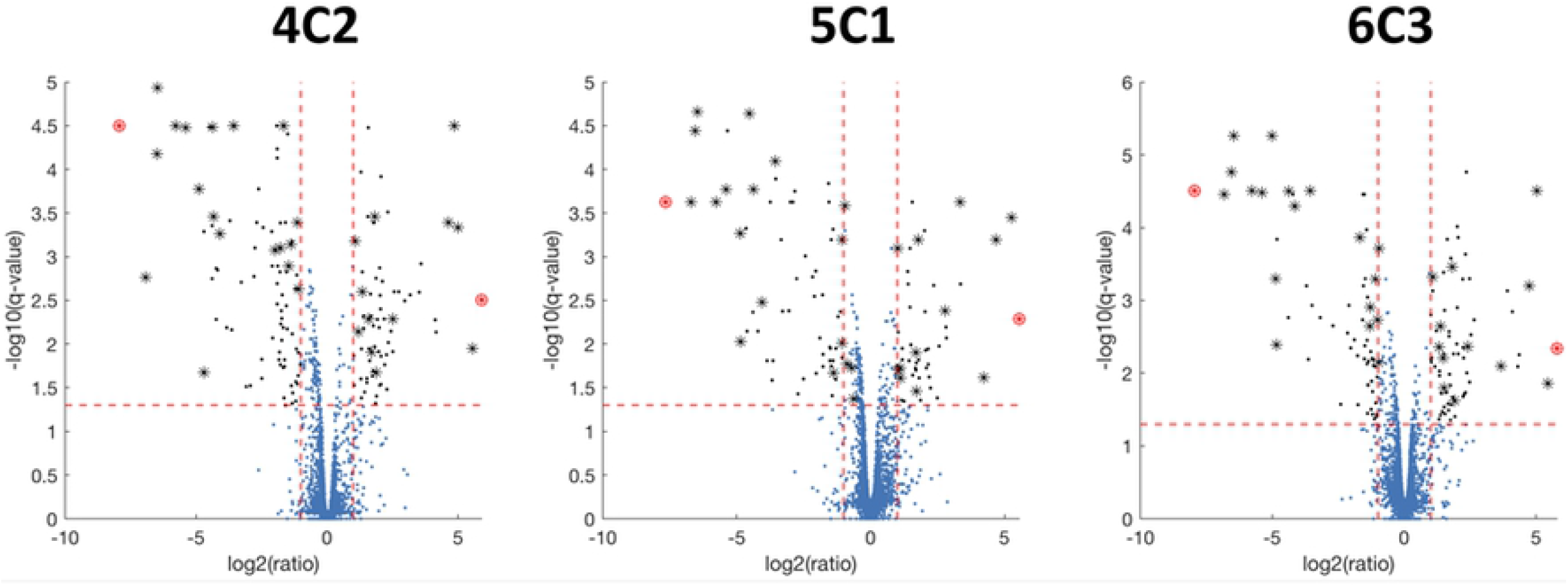
Mitochondrial membrane potential in IL3000 and 4 diminazene-resistant clones, determined by flow cytometry. A. Sample profiles for TMRE fluorescence of 10,000 cells was obtained on a FACSCalibur flow cytometer, using the FL2-height detector. Sample graphs of the IL3000 DA-sensitive parental strain and the 6C1 DA-Res clone are given. B. Average (and SEM, n=3) percentage of cells with a fluorescence >80 A.U. in the analysis for IL3000 and 4 DA-Res clones. Significant differences with the sensitive control are indicated with red stars; difference between 5C1 and the other three DA-res cell lines are indicated with blue stars. ***, P<0.001.

#### Genomic analysis of DA resistance

DA-resistant *T. congolense* were analyzed for SNPs and indels by Illumina paired-end whole genome sequencing. Across the 4 samples (wild type and DA-resistant clones 4C2, 5C1 and 6C3) there were a total of 158,396 SNPs and 28,640 indels (see S1 File and S2 File for the statistics on SNPs and indels, respectively). Of the indels, 20,107 were insertions, whilst the remaining 8,533 were predicted deletions. Both datasets were filtered, firstly to remove mutations occurring in all four samples, as these are clearly not related to drug resistance, and subsequently to obtain lists of high, moderate and low impact (as described in the methods) mutations in protein encoding regions (5,263 SNPs and 541 indels) (S3 Table, worksheets “SNPs-HIGH-MODERATE-LOW” and “Indels-HIGH-MODERATE-LOW”). Once mutations that occurred in genes predicted to encode variant surface glycoproteins (VSGs), expression site-associated genes (ESAGs) and retrotransposon hot spot (RHS) proteins were removed, there were 19 high impact SNPs (S3 Table, worksheet “SNPs-Hi-filtered”) and 192 high impact indels (S3 Table, worksheet “Indels-Hi-filtered”), in addition to 1,693 moderate impact SNPs (S3 Table, worksheet “SNPs-Mod-filtered”) and 42 moderate impact indels (S3 Table, worksheet “Indels-Mod-filtered”) predicted to result in non-synonymous mutations for the former, and in-frame deletions/insertions for the latter. Due to the previously described correlation between resistance to the related compound isometamidium chloride and reduced mitochondrial membrane potential [20], as well as reduced potential observed in the DA-resistant cell lines, data were mined to identify genes that could be involved in maintaining or generating this membrane potential. Two copies of a vacuolar-type Ca^2+^-ATPase (TcIL3000.A.H_000569200 and TcIL3000.A.H_000569400) harbored mis-sense mutations in all three resistant lines (TcIL3000.A.H_000569200: A109T in clones 4C2 and 6C3; TcIL3000.A.H_000569400: G855E in clones 4C2 and 5C1). However, no further candidates were found. Two mis-sense mutations were found in folate transporter TcIL3000.A.H_000597400 (A132S and N131Y, both in clone 4C2 only, both heterozygous) and another two in folate transporter TcIL3000.A.H_000597700 (L122R in 4C2 and F371C in clone 6C3; both heterozygous). No mutations were found in the other folate or pteridine transporters. Mis-sense mutations were also found in several copies of an amino acid transporter (TcIL3000.A.H_000388400, TcIL3000.A.H_000388500 & TcIL3000.A.H_000388600), a putative plasma-membrane choline transporter (TcIL3000.A.H_000774600) and two copies of a mitochondrial chaperonin HSP60 gene (TcIL3000.A.H_000771700 & TcIL3000.A.H_000772600) (S3 Table, worksheet “SNPs-HIGH-MODERATE-LOW”).

#### Transcriptomics analysis of diminazene resistance

To ascertain whether differential gene expression could explain increased levels of resistance to diminazene, three replicates of the wild-type parental line, as well as three replicates of each independent resistant clone, were subjected to transcriptomic analysis by Illumina paired-end sequencing. Data was processed as described in the methods, resulting in transcript abundances for 9,360 *T. congolense* genes (S3 Table, worksheet “RNAseq_all”). Data was filtered to obtain differentially expressed genes relative to wild-type (qval ≤ 0.05), resulting in 274, 247 and 265 genes identified as being significantly differentially expressed in DA-resistant clones 4C2, 5C1 and 6C3, respectively, with 120 of these genes being common to all three clones (Fig 10). Genes were then filtered to remove VSGs, ESAGs and RHS proteins, resulting in a final list of 61 genes, of which 15 were annotated as hypothetical (S3 Table, worksheet “RNAseq_filtered”).

**Fig 10.**
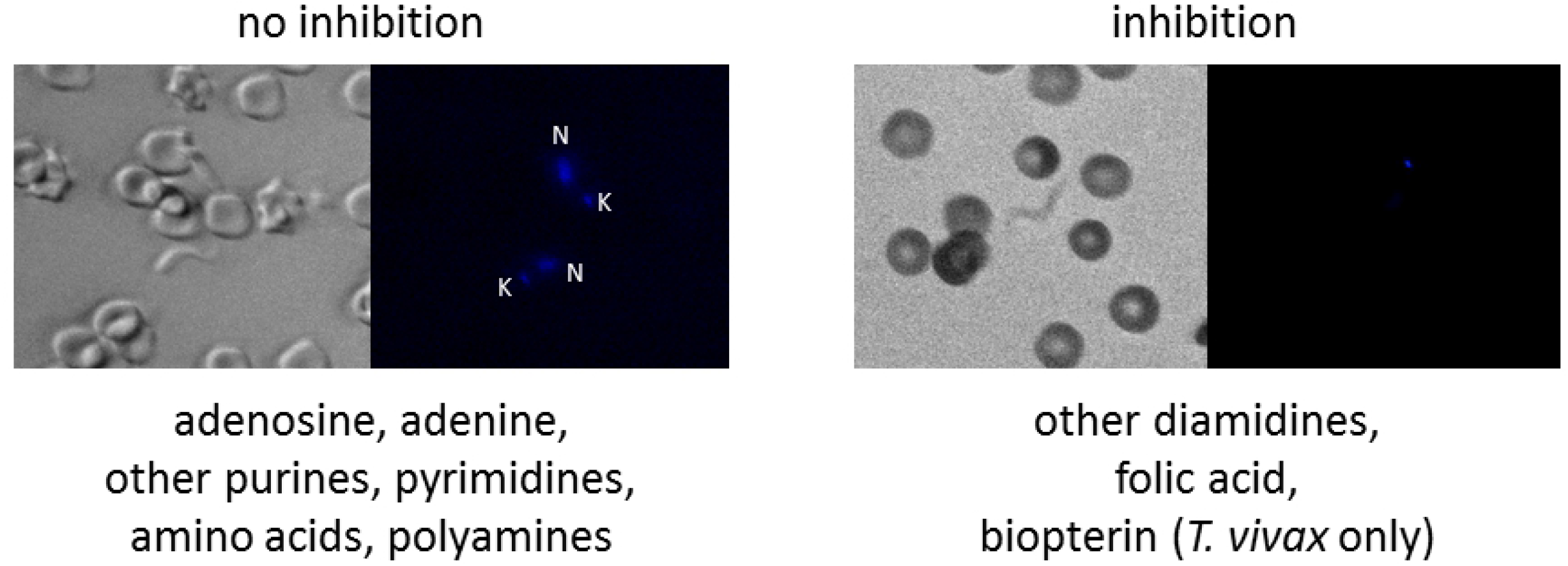
Differential gene expression in three independent DA-resistant clones (4C2, 5C1 and 6C3) of *T. congolense* IL3000. For each DA-resistant clone, Log_2_ fold change was calculated compared to wild-type parasites, as well as significance (q-value), and plotted as a volcano plot. Differential gene expression was deemed significant if the gene exhibited a Log2 fold change of >1 or <-1 in all three clones, as well as a q-value of <0.05 (-Log_10_ q-value of 1.30103) (genes shown as asterisks). The most significantly upregulated (Helicase associated domain (HA2) putative: TcIL3000.A.H_000255300.1) and downregulated (Histone H4 putative: TcIL3000.A.H_000205600.1) genes are shown as red asterisks to illustrate reproducibility across clones.

Among the most significant changes was downregulation of an array of H4 histone encoding genes (mean Log_2_ fold change: −7.842) (S3 Table, worksheet “RNAseq_DOWN”). Furthermore, there was significant downregulation of a putative protein involved in elongation of very long fatty acids (TcIL3000.A.H_000436100.1; mean Log_2_ fold change: - 6.526) and a ZIP Zinc transporter (TcIL3000.A.H_000205900.1; mean Log_2_ fold change: - 4.099). Other downregulated genes of note included a cysteine peptidase, as well as four hypothetical proteins (TcIL3000.A.H_000205700.1, 000206000.1, 000429100.1 and 000429200.1) for which expression was entirely abolished in all resistant clones. None of the hypothetical proteins were found to contain a recognized domain using the SMART [49], interpro [50] and pfam [51] tools. Not all of these may be equally relevant since the locus of 5 genes from TcIL3000.A.H_000205600.1 to TcIL3000.A.H_000206000.1 were all extremely downregulated and this includes a Histone H4, three hypothetical proteins that are not related to each other, and the Zip Zinc Transporter mentioned above.

There were only 13 significantly differentially expressed genes exhibiting a Log2 fold change of >1.0 in all three resistant clones, and these were identified as primarily RNA helicases, in addition to copies of cysteine peptidase and several hypothetical proteins. As the data for the RNA helicases have different significance and expression values, it is unlikely that the identification of this gene family is due to multimapping as in the Histone H4 genes, in this case leading to a firmer conclusion that all individual genes identified are differentially expressed (S3 Table, worksheet “RNAseq_UP”). In some cases, there was significant differential expression in only two of the DA-resistant clones, for example, a predicted long chain fatty acid-CoA ligase (TcIL3000.A.H_00065500.1; mean q-value: 0.0356; mean Log_2_ fold change: −0.297).

## Discussion

In this manuscript we have studied the phenomenon of diminazene resistance in *T. congolense*. This veterinary parasite does not have an equivalent of the *T. brucei* diminazene transporter TbAT1 [31] and thus it is clear that the *T. brucei* model for diminazene resistance, based on loss of TbAT1 function [11,25], is not applicable to *T. congolense*.

We found that it was possible to induce a substantial level of diminazene resistance in *T. congolense* by slowly increasing the drug concentration in the culture medium. The level of DA resistance was entirely stable in all clones, after months of *in vitro* passage in the absence of drug pressure, and repeated cycles of stabilating and reculturing. Remarkably, all the independently generated DA-resistant clones displayed the same level of resistance as well as cross-resistance patterns and growth rates, which could indicate that they had developed very similar adaptations.

One of the main aims of this study was to establish whether DA resistance in *T. congolense* necessarily leads to cross-resistance to other (potential) AAT drugs. Co-resistance with isometamidium chloride has been reported in the field [52–54] but it is not clear whether this constitutes genuine cross-resistance. This conclusion can be hard to reach in the field as cattle are often treated with both drugs and independently-developed resistance to both drugs is therefore a real possibility.

However, there have been reports of single resistance to diminazene and to isometamidium, and more recently to both drugs [1,32,55]. This clearly shows that multiple-resistance is not automatic, as it seems to be for pentamidine and melarsoprol in *T. brucei* sspp. [56]. Our observation that induced DA resistance in *T. congolense* did not change isometamidium sensitivity in any of the independent clones is certainly a strong confirmation of that hypothesis and to the best of our knowledge the first such evidence from a controlled laboratory study since experiments in 1963 by Frank Hawking with multiple *T. congolense* strains in guinea pigs and mice, which also found no evidence of cross-resistance between DA and phenanthridines [57]. We have also attempted to similarly induce isometamidium resistance in *T. congolense*, but this was not successful *in vitro* although it has been achieved *in vivo* [19]. Another important observation on the nature of DA resistance in *T. congolense* is that it was remarkably stable, which, considering the many reports of DA resistance from all over Africa, has important implications for the future utility of the drug.

Based on the above it has been argued that the further development of diamidine drugs against AAT would be unwelcome and only result in more drug pressure on populations that already harbor DA resistance. The strong cross-resistance with furamidines such as DB75 and DB829, observed both *in vitro* and *in vivo* and with field isolates and *in vitro*-adapted clones, would certainly strengthen this argument. However, the complete lack of cross-resistance with pentamidine and the tested series of bis-benzofuramidines demonstrates that only the closest structural analogues display cross-resistance with DA, and that therefore not all diamidines should be ruled out from consideration for further development against AAT. The caveat, however, would be that any new diamidine drug must be active, at a minimum, against the main species causing nagana in sub-Saharan Africa, including DA-resistant strains of each of those species. Given the dearth of knowledge about DA-resistant *T. vivax*, in particular, this would still constitute a significant hurdle at the moment. In this context, the observation that there is no cross-resistance between DA and oxaboroles, promising new agents in development for HAT and AAT [58–60], is highly significant.

In *T. brucei*, drug resistance is mostly linked to changes in drug accumulation, particularly through mutations in transporter genes [12]. In *T. congolense*, isometamidium resistance has also been linked to reduced accumulation [16,17,19,53], but no drug transporters have been identified in this species. We therefore investigated the uptake of [^3^H]-DA in wild-type IL3000 and found it to be very slow and, apparently, low affinity. This indicates that uptake of diminazene could be limiting to its efficacy, whereas in *T. brucei* sspp uptake of diminazene, the furamidines and pentamidine is very fast, and high affinity [7,25,26,43]. We found no consistent indication that DA resistance in *T. congolense* is linked to reduced cellular accumulation of the drug, as the accumulation rate was similar in the parental and resistant clones.

Several compounds were identified as inhibitors of both [^3^H]-diminazene and fluorescent DB75 uptake in *T. congolense*, particularly pentamidine, propamidine and folate. The latter observation indicates the involvement of folate transporters in diminazene uptake, although high concentrations were required for significant inhibition and even then diminazene uptake was only partially inhibited. Clearly, a so-far unidentified, folate-insensitive uptake mechanism is additionally involved in diminazene uptake. The notion of multiple low-affinity entry routes was further strengthened by the demonstration that inhibition by folate and pentamidine was additive, and yet still incomplete. To further investigate whether *T. congolense* folate transporters may contribute to diminazene uptake, all three then known FTs were cloned and expressed in the diamidine- and melaminophenyl-resistant clone *T. brucei* B48 [8]. The expression of IL3000 transporters FT1 and FT3 appeared to induce a small but significant sensitization to diminazene and its structural analogue DB829, but not to pentamidine or oxaborole AN7973; IL3000 FT2 trended in the same direction as the other two transporters but the effect did not reach statistical significance. The folate transporters of two DA-resistant isolates from Cameroon displayed several SNPs relative to the IL3000 sequences but if anything displayed higher levels of sensitization than the homologues from the sensitive strain, and FT2 from both resistant isolates significantly sensitized to all three diamidines including pentamidine, but not to AN7973. These observations show that these folate transporters have a minor capacity for diamidine transport, but also that SNPs observed in these genes are not the cause of DA resistance, either in the field or in our laboratory-adapted clones. The lack of genuine resistance mutations in the folate transporters is presumably linked to the observation that at least two of those transporters have this capacity, and that the FTs mediate only a proportion of DA uptake, considering the modest effect of 1 mM folate on [^3^H]-diminazene uptake.

Based on the above reasoning we must conclude that diminazene resistance in *T. congolense* is not principally the result of a reduced rate of cellular diminazene accumulation. The alternative cause of resistance would be in changes of the target but in the case of diminazene and other cationic drugs this is the kinetoplast DNA [22,46–48], located in the mitochondrion. While dyskinetoplastic *T. brucei* have been described, including as part of the adaptation to isometamidium [21], and *T. evansi* is considered a dyskinetoplastic *T. brucei* [61], dyskinetoplastic *T. congolense* have not been described, and our DA-resistant clones displayed normal kinetoplasts as observed by DAPI staining. Thus, the DA resistance phenotype cannot easily be attributed to changes in the target either. However, in order to reach and disable the kinetoplast, the drug must accumulate inside the mitochondrion – a process that is dependent on the mitochondrial membrane potential Ψm. We observed, in all four DA-resistant clones investigated, a strong and statistically significant reduction in Ψm, which would serve to diminish the entry of diminazene into the mitochondrion, where the drug would otherwise be strongly accumulated, bound to kDNA. This is particularly important as the entry of the drug into *T. congolense* is low affinity and inefficient, as opposed to the energy-dependent, high affinity process in *T. brucei*. Diminazene uptake in *T. congolense* is thus very likely equilibrative rather than concentrative, implying that a failure of the drug to accumulate in the mitochondrion will eventually result in a reduced cellular uptake as well.

Genomic and transcriptomic analyses of the 4C2, 5C1 and 6C3 DA-resistant clones were performed to further investigate the causes of resistance. For each clonal line several hundred genes were differentially regulated, relative to the parental clone IL3000. After filtering-out inconsistent and functionally irrelevant returns relatively few genes were significantly downregulated (including H4 histones, a cathepsin-like cysteine protease and several hypothetical proteins with no known domains) or upregulated (mainly RNA helicases and a few hypothetical proteins). The genomic sequencing revealed 19 high impact SNPs and 192 high impact indels. In all three DA-resistant clones sequenced, mis-sense mutations were observed in at least one of two copies of a vacuolar-type Ca^2+^-ATPase. SNPs were also observed in several transporter genes, including amino acid transporters, some of the folate transporter genes and a putative choline transporter, as well as mitochondrial HSP60. While these observations provide potential leads for understanding DA-adaptation, this will clearly require extensive further validation. The very high level of DA-induced SNPs and indels is consistent with the DNA-targeting nature of this minor grove binder.

Altogether, we conclude that a clinically relevant level of DA-resistance in *T. congolense* can be induced *in vitro* and is stable, and not primarily due to reduced DA accumulation but instead linked at least in part to a deficient accumulation into the mitochondrion due to reduced mitochondrial membrane potential. DA uptake is low affinity and partially mediated by the *T. congolense* folate transporters, but also by separate transporters that are sensitive to pentamidine. These results constitute the first systematic analysis of DA-transport and resistance mechanisms in *T. congolense* and provide some insights important for the development of new drugs against AAT.

## Materials and Methods

### 1. *In vitro* culture of *T. congolense*

Cultures of bloodstream forms of Savannah-type strain IL3000 and the adapted clones derived from this strain were cultured exactly as described by Coustou et al. [62], in a basal MEM-based medium supplemented with 20% goat serum 14 μL of β-mercaptoethanol, 800 μL of 200 mM glutamine solution, and 10 mL of 100× penicillin/streptomycin solution per liter of medium (pH = 7.3). *T. congolense* were cultured in 6- or 24-well plates at 34 °C and 5% CO_2_. For adaptation to DA, cultures were serially passaged in the highest concentration of DA tolerated, which was increased as the cells adapted, starting at 50 nM DA (Sigma). Thus one 24-well plate contained 1 row of ‘no drug’ control culture of IL3000 bloodstream forms and 3 rows of independent cultures in the presence of DA, in three drug concentrations per passage, one above and one below the concentration under test..

### 2. Resazurin assay with *T. congolense*

This drug susceptibility assay was performed essentially as described [63] and is based on only live cells reducing the blue and non-fluorescent viability indicator dye resazurin sodium salt (‘Alamar blue’; Sigma) to pink, fluorescent metabolite resorufin [36]. The fluorescence was quantified using a FLUOstar Optima (BMG Labtech, Durham, NC, USA) at wavelength of 540 nm (excitation), 590 nm (emission) and the data plotted to a sigmoidal curve with variable slope using Prism 5.0 (GraphPad Software Inc., San Diego, CA, USA). The assay was set up in 96-well plates with the test drug added at doubling dilutions in culture medium, usually starting at 50 μM, over 1 or 2 rows of the plate, with the last well receiving 100 μL culture medium as a drug-free control, i.e. either 11 or 23 doubling dilutions. To each well, 100 μL of culture medium containing 5 × 10^5^ bloodstream *T. congolense* cells was added, and the plate was incubated for 48 h at 34 °C/5% CO_2_, followed by the addition of 20 μL of a 125 mg/mL solution of resazurin sodium salt in phosphate-buffered saline (PBS) and a further incubation period of 24 h.

### 3. *In vivo* experiments and ethics statement

Drug sensitivity testing in mice used the standardized single-dose procedure of Eisler et al. [64]. Groups of 6 NIH mice (Envigo) were used throughout, and infected i.p. with 10^5^ *T. congolense* of strain IL3000 or KONT2/151 in phosphate-buffered saline pH 7.4 (PBS). At 24 h post infection (PI) a single dose of DB829, AN7973 or vehicle (DMSO, drug-free control) was administered i.p., at doses 10, 20 or 40 mg/kg bw for DB829 and 5, 10 or 20 mg/kg bw for AN7973. Parasitemia was monitored daily, scored by microscopic analysis of blood from a tail prick from each infected animal. Any mouse found to have high parasitaemia during the experiment, or show any signs of suffering was euthanized by CO_2_ inhalation. All animal experimentation was performed at the University of Glasgow Joint Research Facility under the supervision of trained professionals; the facility is regularly inspected by a UK Home Office Inspector and adheres to all national and international regulations as stipulated by the UK Home Office for animal care and in accordance with the Animals (Scientific Procedures) Act 1986 as amended in 2012. The procedure had been expressly approved and licensed by the UK Home Office (project license PCF371688) and the experiment was performed by a trained animal technician under his personal Home Office license (PIL601/12386).

### 3. Fluorescence microscopy screen for diamidine uptake inhibitors

NIH mice (Envigo) were infected with 100,000 trypanosomes of either *T. b. brucei* strain Lister 427 (i.p.), *T. congolense* strain IL3000 (i.p.) or *T. vivax* strain Y486 [65] (i.v.) in PBS; each parasite species was injected in sterile saline solution. At peak parasitemia the mouse was sacrificed by CO_2_ inhalation and the blood collected by aortic bleed. This was kept on ice if not being used immediately for an experiment.

Approximately 1 mL of whole blood was spun at 1000 × *g*, and supernatant, buffy coat and a visible layer of parasites were removed to a new tube and spun again. The resulting pellet was washed with 1 mL of CBSS buffer (25 mM HEPES, 120 mM NaCl, 5.4 mM KCl, 0.55 mM CaCl_2_, 0.4 mM MgSO_4_, 5.6 mM Na_2_HPO_4_, 11.1 mM glucose; pH adjusted to 7.4 with NaOH), and the pellet resuspended in CBSS buffer at an appropriate concentration to visualize the parasites (usually 1 mL). The duration of viability of parasites was assessed at room temperature, 34 °C and 4 °C prior to completion of experiments. *T. congolense* parasites performed similarly when kept at 34 °C and room temperature during experiments, therefore experiments were conducted at room temperature; *T. vivax* parasites were motile for longer and took up DB75 at a faster rate at 34 °C, so were kept at this temperature as much as possible during experiments (but not while under the microscope for observation).

For non-inhibited cells 10 μM of DB75 was added and fluorescence observed using a Zeiss Axioscope fluorescence microscope, using the DAPI filter set (λ_exc_ = 330 nm, λ_em_ = 400 nm) as well as brightfield. Images were obtained with Openlab imaging software (Improvision, Coventry, UK). For inhibited cells, the test inhibitor was added to the desired concentration (see S1 Table) just before the addition of 10 μM DB75 and observation of fluorescence as before. Each potential inhibitor was tested at least twice on parasites from different mice, those which inhibited up to 4 times. Every experiment had cells with no inhibitor added visualized in parallel to control for slight variation in timing of appearance of fluorescence in kinetoplasts and nuclei between batches of ex vivo parasites.

### 4. Transport assays

Transport assays for *T. congolense* bloodstream forms were performed exactly as described for *T. b. brucei* bloodstream forms [66] and *Leishmania major* promastigotes [67]. Depending on the assay, parasites were either purified from infected blood using a DEAE-cellulose column and a 6:4 ratio PSG buffer as described [68], or collected from culture. Briefly, cells were washed into assay buffer (33 mM HEPES, 98 mM NaCl, 4.6 mM KCl, 0.55 mM CaCl_2_, 0.07 mM MgSO_4_, 5.8 mM NaH_2_PO_4_, 0.3 mM MgCl_2_, 23 mM NaHCO_3_, 14 mM glucose, pH 7.3) and adjusted to a density of 10^8^/mL just before use. 100 μL of the cell suspension was added to 100 μL of radiolabelled substrate (^3^H-DA or ^3^H-pentamidine) in assay buffer, sometimes also containing a competitive inhibitor at 2× final concentration, atop a layer of oil (1:7 of mineral oil and di-n-butyl phthalate (Sigma)) in a 1.5 mL microfuge tube. After a predetermined incubation time the incubation was stopped by the addition of 1 mL ice-cold ‘stop solution’ (1 mM/250 μM unlabelled permeant in assay buffer) and cells were separated from the extracellular radiolabel by centrifugation through the oil layer (1 min at 13,000 rpm in a microfuge). The cell pellets were harvested into scintillation tubes by cutting off the tip of the microfuge tube after flash-freezing in liquid nitrogen, incubated with 2% SDS for at least 1 h, and overnight with scintillation fluid (Optiphase HiSafe III, Perkin-Elmer, Waltham, MA, USA) before being agitated overnight in scintillation fluid (Optiphase HiSafe III, Perkin-Elmer, Waltham, MA, USA). Tubes were then read using a Hidex 300 SL scintillation counter (Lablogic, Sheffield, UK). Raw disintegrations per minute (dpm) reads were converted to units of pmol(10^7^ cells)^−1^ and corrected for background radiation and non-specific radiolabel association, by subtracting dpm counts from no-label controls and parallel determinations in the presence of saturating levels of non-labelled permeant, respectively. Radiolabelled ring-[^3^H]-DA was custom-made by PerkinElmer (CUST78468000MC; 60.7 Ci/mmol) and [^3^H]-pentamidine isethionate was custom-made by Amersham (TRQ40084; 3.26 TBq/mmol). [3,5,7,9-^3^H]-folic acid (ART 0125; 56.8 Ci/mmol) was from American Radiolabelled Chemicals. Staistical analyses were performed with GraphPad Prism 8.1.

### 5. Cloning and expression of *T. congolense* folate transporters

Three folate transporters were identified in the original *T. congolense* IL3000 genome available in TriTrypDB (version 1 of the genome, released Oct 20 2010). The gene designated as Folate Transporter (FT) 1 (TcIL3000.10.7850) was annotated as a predicted pteridine transporter, and FT2 (TcIL3000.0.06950) and FT3 (TcIL3000.0.13340) were predicted folate transporters.

Subsequent to the completion of the cloning work described in this paper, the *T. congolense* IL3000 2019 genome was released in TriTrypDB (release Nov 6 2019); this contains several (additional) predicted genes all annotated as putative folate transporters. TcIL3000.A.H_000799700 aligns with our FT1 (GenBank accession numbers in Table 2 and S2 Fig). Three tandemly arranged, almost identical genes (97.8-98.4% identity), TcIL3000.A.H_000597400, TcIL3000.A.H_000597500 and TcIL3000.A.H_000597700, align with our FT2 with 97.3-99.8% identity by Clustal Omega analysis. And two genes annotated as being pseudogenes due to missing START and STOP codons, TcIL3000.A.H_000531600:pseudogene and TcIL3000.A.H_000558900:pseudogene, align with our FT3. The predicted CDS of TcIL3000.A.H_000531600 is the same as the IL3000 FT3, only having a predicted N-terminal extension with no START codon. The last 27 amino acids and STOP codon that are missing from the sequence given on the TriTrypDB gene page are actually present when the full contig is examined. There are long stretches of TcIL3000.A.H_000558900 that are divergent (67.9% identical by Clustal Omega analysis). All are aligned in S2 Fig.

Based on the TritrypDB sequences obtained, primers were designed to amplify the three genes with either an *Apa*I or *Hind*III 5’ restriction site and a 3’ *Xba*I restriction site. The Primer sequences were as follows: FT1_F 5’-gggcccATGTTTGGCACGGCTGCG-3’; FT1_R 5’-tctagaCTATCTGCGGGCCTCA-3’; FT2_F 5’-aagcttATGAGCGCACCAACCGAC-3’; FT2_R 5’-tctagaCTACCTTACTTCATTGTCTG-3’; FT3_F 5’-gggcccATGTTTGGCGAAGCAACTG-3’ and FT3_R 5’-tctagaCTACCTTACTTCATTGTCTG-3’. Genomic DNA was extracted from parasites isolated from 1 mL of cardiac puncture blood from mice infected with each strain: IL3000, KONT2/133 and KONT2/151, using a Qiagen DNeasy blood and tissue kit, following the manufacturer’s instructions. Genes were amplified from the relevant gDNA using the primers given above and Phusion polymerase (New England Biolabs), following the manufacturer’s protocol. DNA fragments of approximately the correct size were extracted from 1% agarose gels and ligated into the pGEMT-Easy (Promega) subcloning vector. Positive colonies were identified by screening using the M13F/M13R sequencing primers on the vector, and sequenced by Sanger sequencing (Source BioScience, Nottingham UK). The identified sequences were ligated into the vector pHD1336 and transfected into *T. b. brucei* strain B48 [8] using an Amaxa Nucleofector, program X-01, and TbBSF buffer as described [31,69]. Transfectants were cloned out, by limiting dilution, in standard HMI-11 medium [70] containing 5 μg ml^−1^ blasticidin S and clones screened for correct integration of the cassette by PCR. All clones were maintained in HMI-11 using standard growth conditions, 37 °C and 5% CO_2_.

Drug sensitivity of the *T. brucei* cell lines was assessed using the standard resazurin assay, essentially as described for *T. congolense*, above and as described previously [71]. Briefly, the assay was performed in 96-well plates with 11 doubling test compounds dilutions, highest concentration 10 μM, and the 12^th^ well containing cells in drug-free medium. 2 × 10^4^ bloodstream form trypanosomes were added to each well, and the plates were incubated for 48 h at 37 °C/5% CO_2_ after which 20 μL of resazurin solution was added to each well and the plates incubated for another 24 h.

### 6. Mitochondrial membrane potential (Ψm) assay

Changes in Ψm were determined by flow cytometry using the indicator dye tetramethylrhodamine ethyl ester (TMRE) as the probe, exactly as previously described [47,63] Briefly, bloodstream forms of 1×10^7^ *T. congolense* IL3000 and several IL3000-adapted DA-Res clonal lines in mid-log growth phase were centrifuged at 1600 × *g* for 10 min at 4 °C and washed in 1 mL pf PBS (pH 7.4) and finally resuspended in 1 mL PBS containing 200 nM TMRE. These cells were incubated for 30 min at room temperature and then placed on ice for a further 30 min prior to analysis on a Becton Dickinson FACSCalibur™ system (BD Biosciences) using a FL2-height detector, and CellQuest and FlowJo 10 software.

### 7. DAPI staining of nuclei and kinetoplasts

Approximately 1 × 10^6^ cells were harvested in Eppendorf tube and centrifuged at 2,400 rpm for 10 min in a microfuge. The supernatant was discarded, the cells were resuspended in 1× PBS and washed by centrifugation (2,400 rpm for 5 min). The supernatant was discarded, the cells were resuspended in 50 μL 1× PBS pH 7.4, spread out on a microscope slide and allowed to dry for 30 min. The cells were then fixed by flooding the slide with 4% formaldehyde for 10 min, followed by washing twice in 1× PBS. The slides were allowed to air dry before placing a drop of Vectashield mounting medium containing DAPI (4’,6-Diamidino-2-phenylindole dihydrochloride) (Vector Laboratories) on them. The slides were then covered with a cover slip and the edges sealed with nail varnish. The slides were viewed under a DeltaVision microscope (GE Healthcare) using softWoRx software and the images were processed using ImageJ software.

### 8. RNAseq

Total RNA isolations were performed on 1 × 10^7^ cells of a parental strain of wild-type *T. congolense* IL3000, as well as 3 replicates of each of the three independent DA-resistant clones (4C2, 5C1 and 6C3), using a commercial kit (RNeasy, QIAgen). Samples were enriched for mRNA by Poly-A selection, using the TruSeq stranded mRNA sample preparation kit (Illumina, catalogue number 20020594) and subsequently quantified using Qubit high sensitivity reagents (Thermo). Average library size was determined using DNA high sensitivity reagents for the Agilent bioanalyser 2100 (Agilent). For sequencing, 2×75 bp paired-end sequencing was carried out using an Illumina NextSeq 500 system (Illumina), resulting in an average of 12 million reads per sample. For RNA-seq data processing, reads contained in the resultant fastq files were aligned to the PacBio genome of the *T. congolense* strain IL3000 [72] and gene abundances quantified, in both cases using Kallisto with default parameters (-b 100, -t 8) [73]. Statistical analyses of differential expression between the wildtype parental strain and each of the three resistant clones were carried out using the R-based package Sleuth, with default parameters (for each transcript, a second fit was performed to a reduced model that presumes abundances are equal in the two conditions - wild-type and DA-resistant) as outlined in the manual [73], and figures were generated with Matlab (The MathWorks, Inc.). Genes exhibiting significant (q-value <= 0.05) differential expression between wild-type and resistant parasites were tabulated and filtered to remove VSGs, ESAGs and retrotransposon hot spot proteins, and subsequently sorted to obtain genes with significance based on q-value (i.e. p-value adjusted for multiple testing by false discovery rate). The full Sleuth output is available in S3_Table, worksheet “RNAseq_all”).

### 9. Whole genome analysis

For whole genome analysis, DNA was extracted from *in vitro* cultures (3×10^6^ cells) of the *T. congolense* IL3000 wild type line, as well as the 3 DA-resistant lines (4C2, 5C1 and 6C3), using the QIAgen DNeasy kit. Libraries were prepared using the QIAseq FX DNA library kit (QIAgen, cat: 180475). The libraries were quantified using Qubit high sensitivity reagents (Thermo), and average library size was determined using DNA high sensitivity reagents for the Agilent bioanalyser 2100. The four samples were subsequently paired-end (2×75 bp) sequenced using Illumina NextSeq500 to an average of 15 million reads per sample. SNP and indel analysis were carried out on the resulting data using the GATK pipeline [74]. Reads were first trimmed using cutadapt (parameters: -q 30,30 --minimum-length=70 --pair-filter=any) [75] and subsequently aligned to the PacBio *T. congolense* IL3000 assembly (Abbas et al., 2018) using STAR (parameters: --outSAMtype BAM unsorted --readNameSeparator _ --readFilesCommand zcat --runThreadN 8) [76]. The resulting bam files were further processed with Picard (functions: AddOrReplaceGroups & MarkDuplicates) [74] and GATK (functions: SplitNCigarReads, HaplotypeCaller, VariantFiltration, SelectVariants) to generate separate files for SNPs and indels. Further filtering was carried out on these files using the SnpEff tool (parameter: SnpSift.jar filter “(FILTER = ‘PASS’)”) [77]. Final datasets for SNPs and indels were created using the SnpEff command (SnpEff.jar) and data was extracted into tabular form using the SnpSift command [78] (extractFields - CHROM POS ID REF ALT FILTER “ANN[*].GENE” “ANN[*].GENEID” “ANN[*].FEATURE” “ANN[*].EFFECT” “ANN[*].IMPACT” “ANN[*].CODON” “ANN[*].AA” “ANN[*].ALLELE” “GEN[*].GT”). The resulting data was filtered, firstly to remove SNPs occurring in both wild-type and resistant lines, and subsequently to select for SNPs and indels predicted to have a high (a variant that is assumed to result in a highly disruptive impact on the protein such as truncation or loss of function), moderate (non-disruptive variant that is assumed to impact protein effectiveness such as non-synonymous SNPs or in frame deletions) or low (Mostly harmless variants such as synonymous SNPs) impact, in both cases using Microsoft Excel.

Sequence data has been deposited in the European Nucleotide Archive (accession number PRJEB39051).

## Acknowledgements

The Global Alliance for Livestock Veterinary Medicines (GALVmed) is gratefully acknowledged for funding this study. We thank professor Dave Boykin (Georgia State University, Atlanta, GA, US) and professor Richard Tidwell (University of North Carolina at Chapel Hill, Chapel Hill, NC, USA) contributed diamidine compounds, Professor Jan Van den Abbeele for the *T. congolense* KONT2/133 and KONT2/151 isolates, and Pfizer, Anacor, and the Bill and Melinda Gates Foundation for the donation of oxaborole compounds. Mr. Craig Lapsley is gratefully acknowledged for making the sequencing libraries.

## Supporting information

S1 Fig. Adaptation of TcoIL3000 to diminazene aceturate *in vitro*.

S2 Fig. Amino acid alignments of *T. congolense* folate transporters.

S3 Fig. Fluorescence micrographs after DAPI staining of *T. congolense* cells, showing nucleus and kinetoplast.

S1 Table. Potential inhibitors tested and effects on appearance of DB75 fluorescence in kinetoplast and nucleus of *T. congolense* and *T. vivax*.

S2 Table. Identification of amino acid differences in folate transporters of three different *T. congolense* strains.

S3 Table. RNAseq and genomic sequence data of *T. congolense* IL3000 and DA-Res

S1 File. Statistical analysis of SNPs in the *T. congolense* DA-Res genomes

S2 File. Statistical analysis of indels in the *T. congolense* DA-Res genomes

